# Diet- and metabolic state-dependent remodeling of the mouse brain lipidome

**DOI:** 10.64898/2026.02.27.706713

**Authors:** Adélaïde Bernard, Kevin Huynh, Jen Xin Fach, Hang Yi Woo, He Liu, Yingying Liu, Natalie Mellet, Peter Meikle, Brian G Drew, Yi Wang

**Author notes:** These authors contributed equally (Brian G Drew, Yi Wang).

## Abstract

The hypothalamus and brainstem are key hubs of metabolic control that undergo dynamic molecular adaptations in response to changes in energy availability. This remodeling is associated with changes in the expression and activity of enzymes linked to energy homeostasis but also importantly, lipid metabolism. Given that lipids account for ∼50% of the brain’s dry weight, it is likely that lipid metabolism is a major determinant of brain function. Therefore, understanding how the hypothalamic and brainstem lipidome adapts to metabolic perturbation is key to understanding tissue function and metabolic health. Here we characterize the remodeling of ∼750 lipid species in mouse hypothalamus and brainstem, as well as the cerebrospinal fluid and plasma, in response to a metabolic challenge (an *Ad Libitum*-*Fasting*-*Refeeding* cycle). We show that around 45% and 36% of lipids in the hypothalamus and brainstem respectively, exhibit reversible, nutritional state-dependent remodeling during this metabolic challenge, and that this remodeling is substantially impacted by long-term high fat diet intervention. Of note, targeted analysis of specific lipids revealed that certain fatty acids were affected by this intervention in the hypothalamus and brainstem, most strikingly defined by the reversible fasting-induced increase in linoleic acid (18:2)-containing phosphatidylcholines in both the hypothalamus and brainstem, an effect that is abolished by high fat diet intervention. Such precise and intervention-specific regulation of linoleic acid (18:2)-containing phosphatidylcholines provides a previously unrecognized role for this lipid in the physiological response to fasting. Thus, these findings demonstrate that the brain lipidome undergoes robust, nutritional state-dependent remodeling, and provide a comprehensive resource for investigating its role in regulating metabolic adaptations.

## Introduction

The brain has the second highest lipid content in the mammalian body behind adipose tissue. Lipids have several important functions including being structural elements of cell membranes and myelin, which are essential for both inter- and intra-neuronal communication. Lipids are also involved in the regulation of cellular functions such as signaling, apoptosis, mitochondrial function, immune function, and metabolism^1^. As an energy source, lipids such as triglycerides (TGs) stored in lipid droplets, are hydrolyzed into fatty acids which are converted into fatty acyl-CoA and then transported by acylcarnitine into the mitochondria. Inside mitochondria, fatty acids are released from carnitine which can undergo β-oxidation and ultimately ATP generation^2^.

β-oxidation is widely used in metabolically demanding tissues such as liver, heart, and skeletal muscle as a means of generating energy. On the contrary, the brain has been previously thought to rely almost solely on glucose for energy production - without storage and dissipation of lipids as fuel^3^. This is interesting considering that the brain is the most energy-demanding organ in the human body, using ∼20% of the body’s total energy, with half of its dry weight comprising lipids.

Most recent discoveries have shown that specific cells in the brain, such as glia cells, are indeed able to utilize fatty acids as energy source^4^ or reserve^5^. Moreover, fatty acid oxidation has been shown in astrocytes to contribute to about 20% of the brain’s total energy needs^4^, whilst in oligodendrocytes it has been proposed to serve as energy reserves to support mouse optic nerve during glucose deprivation^5^. Importantly, multiple recent studies have demonstrated that neurons are able to metabolize fatty acids through β-oxidation to maintain neuronal energy and function, but specifically when glucose is limited^6-9^. Together, these studies provide new and compelling evidence that lipid-derived fatty acids serve as an alternative energy source to sustain brain function during metabolic challenges.

Further evidence for the use of lipids as an energy source in the brain has emerged when studying myelin, which is primarily composed of lipids, and is synthesized by oligodendrocytes to surround and encase axons. This study demonstrated that brain myelin content in humans can be temporarily and reversibly diminished by prolonged endurance exercise, a model of energy depletion, suggesting that myelin lipids may act as energy reserves in extreme metabolic conditions^10^. The above findings collectively indicate that the lipidome, although a major structural component of the brain, might be more malleable to metabolic challenges than was previously anticipated^11^.

The hypothalamus and brainstem are the two major brain regions that coordinate dynamic adaptations to metabolic challenges. Lipid sensing is a crucial aspect of the regulation of energy balance in the hypothalamus and brainstem, as both these regions express receptors that can detect lipids in the extracellular environment, such as free fatty acids^12^. Moreover, it has been shown that enzymes involved in lipid metabolism, such as plasticity-related gene-1 (PRG-1)^13^, fatty acid synthase (FAS)^14,15^, and sterol regulatory element-binding protein-1c (SREBP-1c)^16,17^, play roles in the regulation of energy balance through modulating intracellular lipid synthesis and conversion in the hypothalamus. Those lipids synthesized or converted in the hypothalamus can mediate neuronal activity and synaptic signaling to modulate feeding behavior and energy expenditure. However, very little is known about how the lipidome in the hypothalamus and brainstem adapt to metabolic challenges. Here, we report the changes of ∼750 individual lipid species from 45 lipid classes and subclasses in mouse hypothalamus, brainstem, cerebrospinal fluid and plasma to an *Ad Libitum–Fasting–Refeeding* cycle, as well as the impact of high fat diet interventions on these adaptations.

## Results

To understand how the lipidome in the hypothalamus and brainstem adapt to metabolic challenges, as well as the impact of high fat diet interventions on these adaptations, we performed experiments in male C57BL/6 mice that were fed chow, high fat diet for 3 days (HFD 3d), or HFD for 8 weeks (HFD 8w, Fig. 1A&B). These mice were then subjected to a “metabolic challenge” consisting of an *Ad Libitum*-*Fasting*-*Refeeding* (AL-F-R) cycle^21^. We chose these particular feeding protocols of HFD, because it is known that consumption of HFD can induce maladaptation such as leptin resistance in the hypothalamus very rapidly, within 24-48 hours, when excessive weight gain has not been developed^18^. Moreover, 8 weeks of HFD feeding is enough to induce obesity and whole-body metabolic dysfunction in C57BL/6 mice^19,20^. Therefore, these timepoints were chosen to allow us to study the impact of HFD on the remodeling of brain lipidome to metabolic challenges both on a short term, before the onset of obesity, and on a long term, after full development of obesity and metabolic dysfunction.

**Figure 1:**
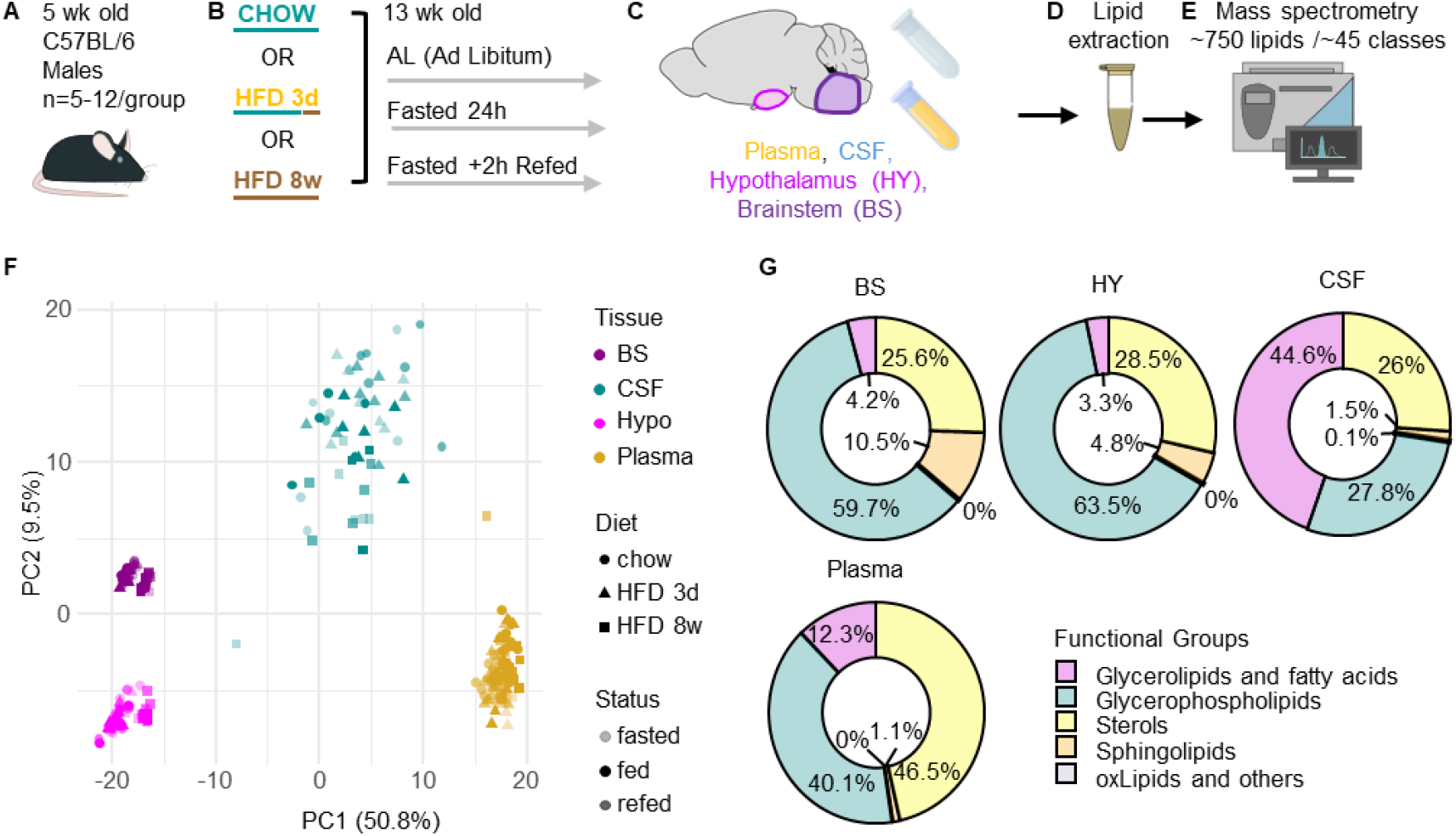
Brain, CSF and plasma have different lipid composition. Experimental design (**A-E**): C57BL/6 male mice (n=5-12/group, **A**) were either fed standard chow or 3d or 8w of HFD, and either fed *ad libitum* (AL), fasted for 24h (F) or fasted for 24h and refed for 2h(R) (**B**) before tissue collection at 13 weeks of age. Plasma, Cerebrospinal Fluid (CSF), Hypothalamus (HY) and brainstem (BS) were collected (C). Lipid were extracted (D), separated using liquid chromatography and quantified mass spectrometry (E). **F** PCA analysis done on 511 common lipid species from n= 267 samples. **G** Proportions of lipids belonging to different functional groups of lipids in plasma, cerebrospinal fluid (CSF), brainstem (BS) and hypothalamus (HY) of mice fed chow.

Within each diet group, mice were randomly allocated to *Ad libitum* (AL), fasted for 24 hours (F), or fasted for 24 hours and refed for 2 hours (R), to measure their responses to an AL-F-R cycle (Fig. 1B). Body and tissue weights of these mice were provided in Suppl Fig. 1.

### Lipid composition in brain tissues is distinct from plasma and CSF

We collected the hypothalamus (HY) and brainstem (BS, specifically Pons and Medulla Oblongata) from these mice, and then extracted and analyzed the lipid composition in those tissues using mass spectrometry (Fig. 1C&D). Cerebrospinal fluid (CSF) and plasma were also collected and analyzed to understand the interactions between brains tissues, CSF and plasma (Fig. 1C&D). A principal-component analysis (PCA) demonstrated clear separation of the various sample types based on lipid profiles (Fig. 1F).

Upon demonstrating that there were distinct and varied lipidome landscapes in these tissues, we sought to investigate what characterized these differences, and thus we plotted the major classes of lipids in doughnut charts for the 4 different tissues (Fig. 1G). In mice fed *Ad libitum* chow, the HY and BS contained around 60% glycerophospholipids, the major component of cell membranes, followed by 25-28% sterols, 5-10% sphingolipids, and less than 5% glycerolipids and fatty acids. In contrast, the plasma and CSF contained higher proportions of glycerolipids and fatty acids (12% and 45% respectively), and lower proportions of glycerophospholipids and sphingolipids, compared to the HY and BS (Fig. 1G). A comprehensive analysis of the proportion of lipid classes within the 5 main functional groups across tissues in *Ad libitum* chow-fed mice was provided in Suppl Fig. 2.

Next, we sought to compare the difference in raw concentrations in lipid classes rather than proportions. Because plasma and CSF are body fluids, analyses were performed within physiologically comparable compartments (i.e. CSF vs plasma; HY vs BS). It is known that the total lipid abundance of human CSF is around 0.2% of plasma levels^22^. Consistent with this observation, our data demonstrated that the overall lipid abundance in plasma was indeed much higher than CSF (normalized to volume) in *Ad libitum* chow mice, except for a few classes of lipids with low abundance in both plasma and CSF, such as phosphatidylserine (PS), lysophosphatidylcholine plasmalogen (LPC P), and phosphatidic acids (PA) (Fig. 2A).

**Figure 2:**
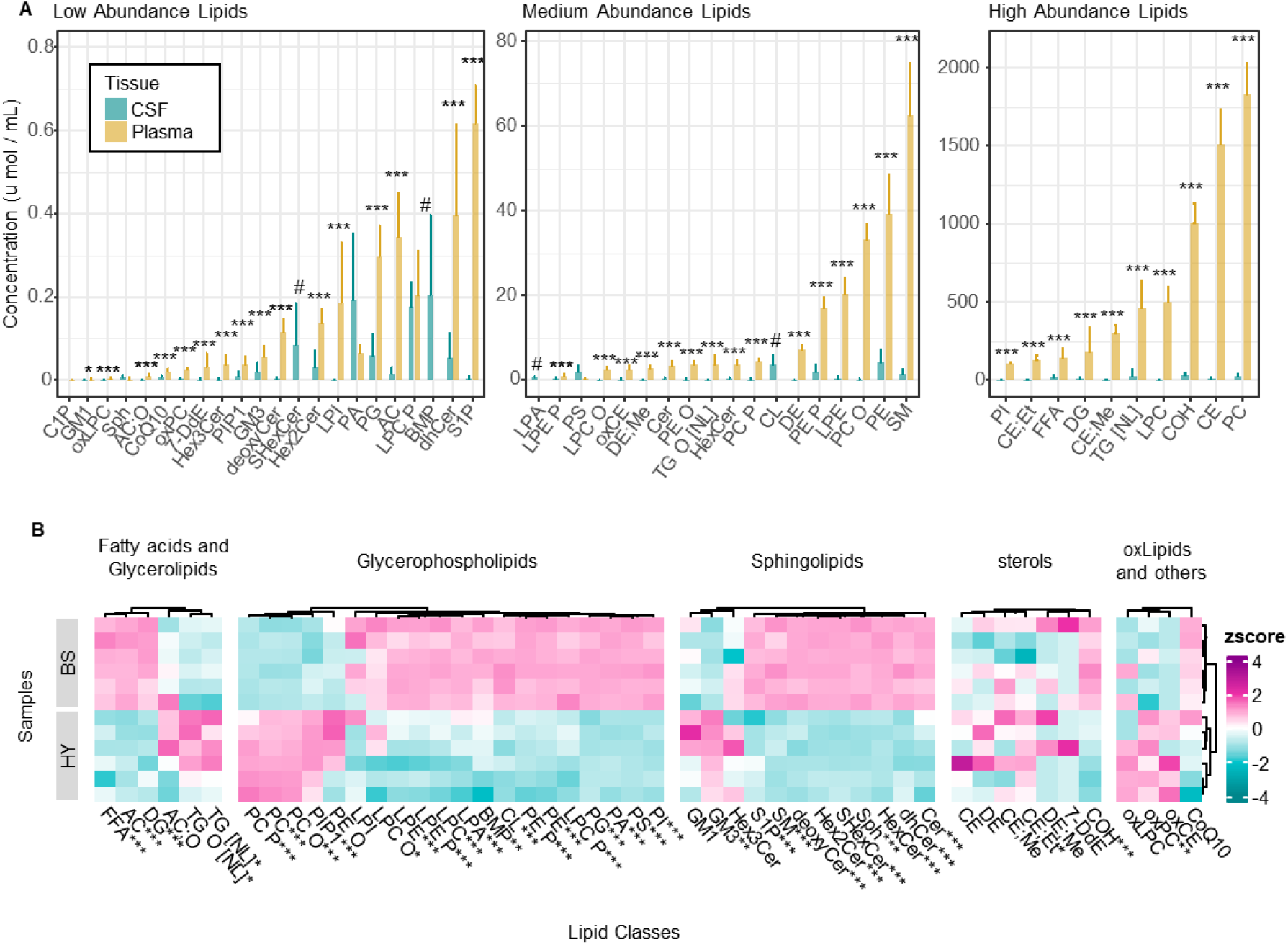
Comparison of lipid classes distribution across tissues. **A** Total concentrations of measured lipid classes in plasma and CSF (umol/mL) **B** Scaled total concentrations of measured lipid classes in brainstem (BS) and hypothalamus (HY). (totals calculated on data normalized to PC+PE+SM) Statistical tests: Student t-test +BH correction. *0.05’ **0.01 ***0.001 #= not measured in plasma. N= 5-7 mice/tissue.

Along the same lines of investigation, we also compared the two tissue regions against each other. The most striking difference from this comparison was that the BS contained higher abundances of sphingolipids. This is likely be explained by the more abundant myelin content in the BS than the HY (Fig. 2B), as sphingomyelins and ceramides are important components of myelin sheath. Of note, the HY contained a significantly higher abundance of PC compared to the BS, while most other glycerophospholipids were more abundant in BS (Fig. 2B).

### Lipidome in plasma, CSF and brain tissues is elastic to an AL-F-R cycle in chow-fed mice

To quantify the adaptations in the lipidome to our metabolic challenge of an AL-F-R cycle, we adopted the concept of “metabolic elasticity”, defined as “the ability of an organism to respond to a disturbance in energy balance and return to its baseline metabolic homeostasis”^21^. It has been shown that a lot of metabolic parameters such as body weight, plasma glucose, insulin, and free fatty acids, are elastic, as they adapt to the disturbance induced by fasting, and subsequently return to baseline after refeeding^21^. To validate this concept in our study, we first measured the body weight, as well as the weights of several peripheral metabolic tissues in response to the AL-F-R cycle, including liver, epididymal fat, subcutaneous fat, and interscapular brown adipose tissue. Our data showed that in chow-fed mice, body weight and the weight of all measured tissues are elastic, as shown by their reduction with fasting, and subsequent rebound towards restoration with refeeding during an AL-F-R cycle (Suppl Fig. 1A-E).

With regards to the lipidome elasticity, we adapted an elasticity scoring system (Gene Elasticity Score, GElaS) that was previously established and validated for transcriptome data, which integrates the change induced by the disturbance, the reversibility of the change once the disturbance ceases, and the statistical significance of those changes^21^. We applied this scoring system to our lipidome data to generate Lipid Elasticity Scores (LElaS). In the GElaS system (gene elasticity), all elasticity scores were positive values, which makes it impossible to distinguish between readouts with an up-down pattern (A-shape) or a down-up pattern (V-shape). In our LElaS system, we gave A-shape lipids a positive, and V-shape lipids a negative elasticity score (Fig. 3A-B).

**Figure 3:**
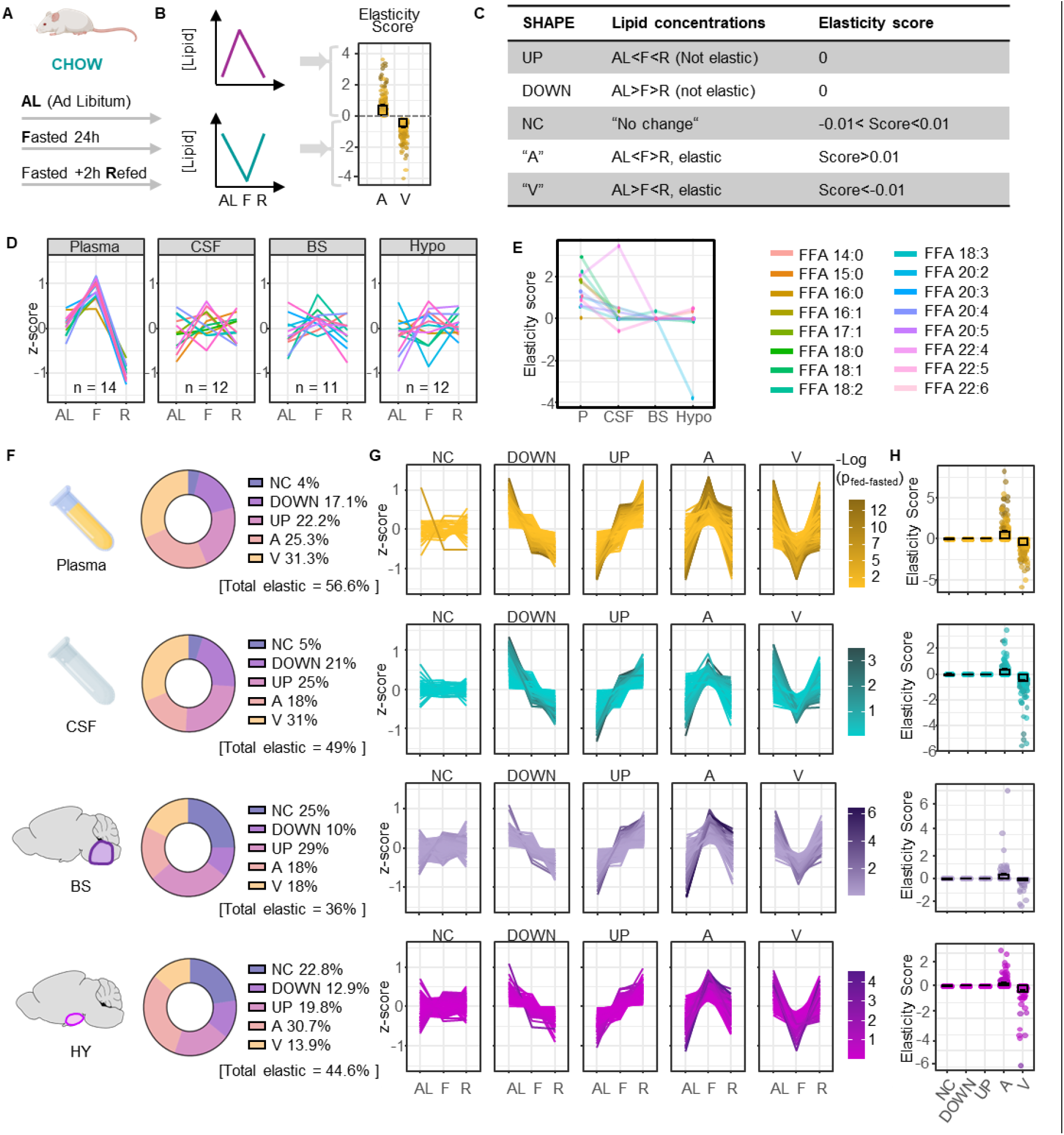
Elasticity of lipid species in tissues from mice fed chow. **A** Experimental design : mice were either fed ad libitum (AL), fasted (F) for 24h of fasted and refed for 2h (R). **B** the changes in lipid species concentrations from AL to F and F to R were compared using ANOVA followed by post-hoc multiple comparisons, and the fold changes and p-values were used to compute elasticity scores for individual lipid species (**B**). Lipid species with an elasticity score lower than 0.01 were categorized as “not changed” (NC) and lipids decreasing with fasting were assigned a negative elasticity score (**C**). **D** Scaled average concentrations of quantified free fatty acids (FFA) on plasma, CSF, BS and HY (left) and their corresponding elasticity score (E). **F-H** Proportion of lipid species assigned to each shape group (NC, DOWN, UP, A, V) in each tissue (left), z-scored concentrations of the corresponding lipid species across feeding status illustrating their changes over the fasting–refeeding cycle (middle), and corresponding elasticity scores for these lipids (right). NC = no change. Colour scale represent –log10(p-value fasted vs fed).

We tested this approach by verifying that lipids show 5 distinct patterns of regulation: consistently increased (Up), consistently decreased (Down), no Change (NC), an up-down pattern (A), or a down-up pattern (V). Those that are Up, or Down, were assigned an elasticity score of 0 (no elasticity), while lipids with a |LElaS| >0.01 were deemed elastic, representing an “A” or “V” elasticity pattern (Fig. 3B-C). We then empirically validated this scoring system by observing the modulation of free fatty acids (FFAs) to an AL-F-R cycle (Fig. 3D). It is known that fasting promotes adipose tissue lipolysis to release FFAs into plasma as an energy source, thus increasing the circulating level of FFAs^23^. Consistent with the literature, all plasma FFAs in our dataset exhibited an “A” shape pattern and were assigned a positive LElaS score (Fig. 3E). Across tissues, the elasticity of the different FFAs did not show consistent patterns with each other, with plasma being the only tissue compartment to demonstrate an overall strong elasticity in FFAs, suggesting each FFA is differentially regulated in different tissues during an AL-F-R cycle (Fig. 3D&E).

We represented the proportion of lipid species that were assigned to 5 pattern groups, in all 4 tissues (Fig. 3F), the scaled concentrations of the corresponding lipid species across feeding states illustrating their changes over the AL-F-R cycle (Fig. 3G), as well as the corresponding elasticity scores for these lipids (Fig. 3H). As expected, the plasma lipidome was the most elastic among the 4 compartments studied, demonstrated by the highest amount of elastic lipid species (56.6% A-shape + V-shape = total elasticity, Fig. 3F). Of note, 44.6% lipids in the HY and 36% lipids in the BS displayed elastic behavior (total elasticity) (Fig. 3F and Suppl Fig. 3A), indicating a significant proportion of brain lipids indeed responded to fasting-induced energy deprivation, and that this response could be returned to baseline after refeeding. To investigate reasons that might explain the higher proportion of elastic lipids in the HY compared to BS, we first compared the total number of lipid species detected in these two brain regions, and confirmed that they contained a similar number of lipid species in all 5 functional groups (Suppl Fig. 3B) Next, To understand which functional group drove the difference between the HY and BS, we compared the number of elastic lipids in different functional groups (Suppl Fig. 3C). Our analysis showed a higher number of glycerophospholipids were elastic in the HY compared to BS, while the number of elastic lipids in other functional groups were similar amongst these two tissues (Suppl Fig. 3C).

Furthermore, when analyzing the top 20 most elastic lipid species among the four different tissues, we identified very different patterns in plasma compared to brain tissues as well as CSF. In plasma, 16 out of the top 20 most elastic lipids were glycerolipids, which typically serve as an energy source (Suppl Fig. 3D-E, Suppl Fig. 4). In contrast, more than half of the top 20 elastic lipids in the HY and BS were glycerophospholipids, that are major constituents of cell membranes and play important roles in various cellular functions including signal transduction, vesicle trafficking, and membrane fluidity (Suppl Fig. 3D-E, Suppl Fig. 4). Thus, we were interested in why the glycerophospholipids might be changing so distinctly in these tissues, and performed further analysis into this phenomenon.

### Long-term high fat diet modulates the elasticity of glycerophospholipids in the hypothalamus and brainstem

It has been reported that both metabolic and gene elasticity are affected by the development of HFD-induced obesity^21^. Therefore, we first compared metabolic elasticity in mice fed on short-term (3 days, before the onset of obesity) and long-term (8 weeks, with obesity) HFD, to chow-fed mice, by comparing their body and tissue weights change during an AL-F-R cycle (Suppl Fig. 1). Our data showed that the metabolic elasticity of both body and tissue weights was progressively abolished by HFD interventions (Suppl Fig. 1F-O).

To understand whether and how HFD intervention modulates lipidome elasticity of the HY and BS, we calculated LElaS scores in mice fed on HFD 3d and HFD 8w (Suppl Fig. 5&6). We observed a change of LElaS patterns in all tissues. Particularly, in the HY and BS, we observed a decrease in elasticity score in mice fed on long-term HFD (Fig. 4A&B). This was characterized by a high proportion of lipids on chow and short-term HFD that behaved in the A-shape pattern and had positive LElaS scores, which then changed to the V-shape pattern with negative LElaS scores on long-term HFD (Fig. 4C&D).

**Figure 4:**
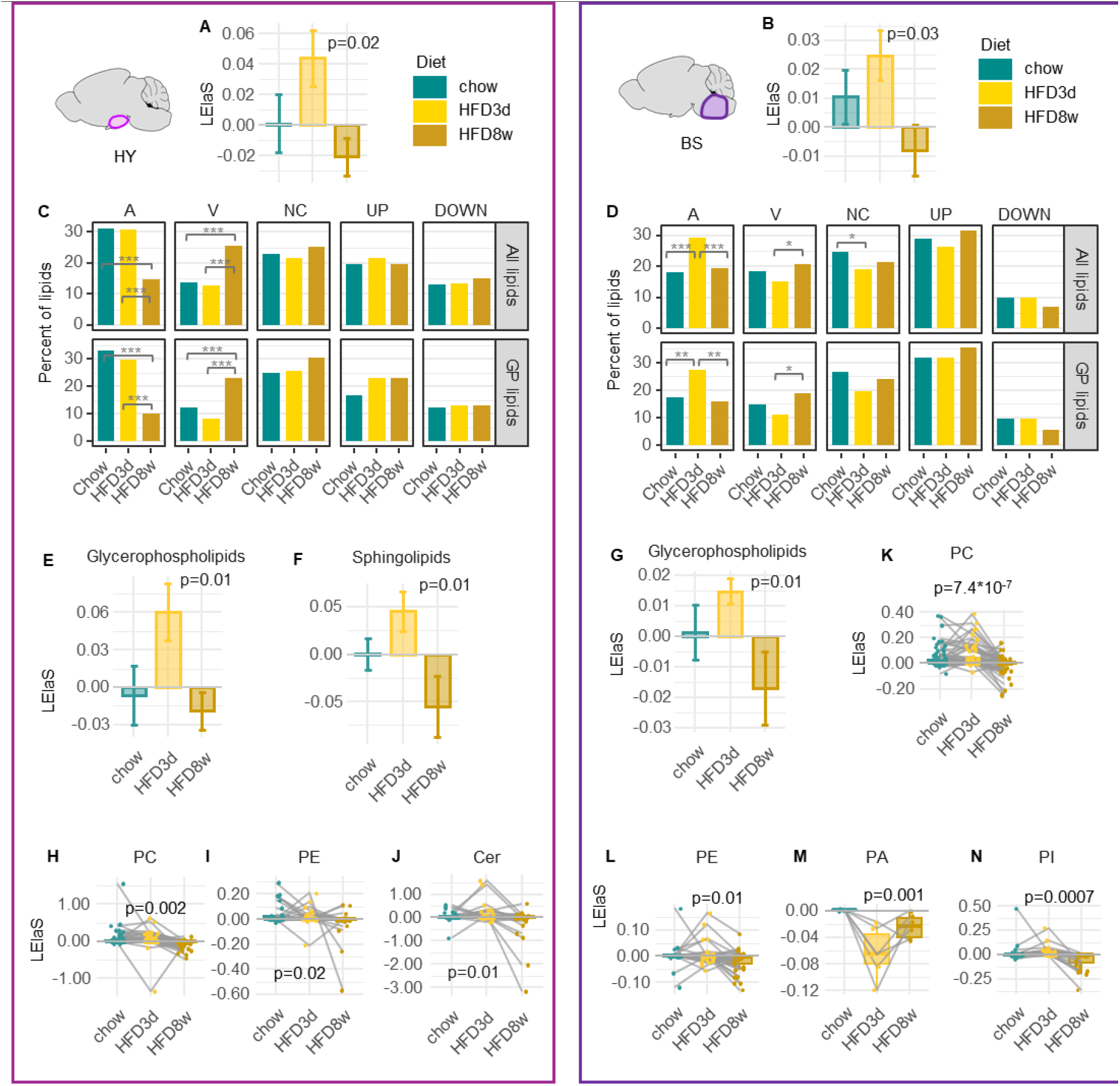
Effect of HFD on the elasticity of lipids in Hypothalamus and brain stem. **A-B** Average signed elasticity scores for all lipids from the brain from mice fed chow, HFD 3d or HFD 8w, in the hypothalamus (A) and the Brainstem (B). **C-D** Proportion (in %) of total number of lipid species detected (all, top or glycerophospholipids only, bottom) that belong to each shape group (A, V, NC (“no change”), UP, DOWN), in the hypothalamus (HY, C) and the brainstem (BS, D) **E-G** Average signed elasticity scores for all glycerophospholipids (E,G) or sphingolipids (F) from the brain from mice fed chow, HFD 3d or HFD 8w, in the hypothalamus (E,F) and the Brainstem (G). **K-N** Signed elasticity scores of lipids belonging to different glycerophospholipid classes that are affected by high-fat diet: Phosphatidylcholine (PC), Phosphatidyl ethanolamine (PE), Phosphatidic acid (PA) and Phosphatidyl inositol (PI) in the hypothalamus (G-H) and the brainstem (I-L). Statistics: Non-adjusted one-way ANOVA p-values comparing diets for each class are displayed on the graphs (A,B,E-N). C-D: For each tissue, differences in the distribution of lipid-shape categories (A, V, NC, DOWN, UP) across diets were tested using a chi-square test of independence on lipid-species counts (diet × shape). Where indicated, post-hoc pairwise comparisons for each shape between diets were performed using two-proportion tests, with Benjamini–Hochberg correction for multiple testing. p=*0.05’**0.01 ***0.001 .

We further performed the same test in different functional groups and classes of lipids. Our analysis showed that it was the glycerophospholipids and sphingolipids in the HY (Fig. 4E&F), and the glycerophospholipids only in the BS (Fig. 4G) that primarily drove this change. In the HY, it was mainly due to the decreased LElaS scores in certain classes of glycerophospholipids including PCs (Fig. 4H) and PEs (Fig. 4I), as well as ceramides (Fig. 4J) in mice fed on long-term HFD, as depicted by a lower overall LElaS at HFD 8w than at chow for these lipid groups (more V-shaped curves result in a lower overall LElaS). In the BS, more glycerophospholipids behaved in a similar way including PCs (Fig. 4K), PEs (Fig. 4L), phosphatidic acids (PAs, Fig. 4M), and phosphatidylinositols (PIs, Fig. 4N). HFD interventions impacted LelaS differently in plasma and CSF, and these results are provided in Suppl Fig. 7.

### Fatty acid chain composition in the hypothalamus and brainstem is impacted by an AL-F-R cycle and high fat diet intervention

Our analysis demonstrated that the elastic portion of the lipidome in the HY and BS is mostly comprised of glycerophospholipids. A major drive of the diversity of glycerophospholipid is the composition of fatty acid chains, which typically vary from 16 to 24 carbon atoms in length and contain zero to six double bonds. Fatty acid chains play important roles in cellular metabolism, including serving as building blocks for cell membranes, providing a major source of energy storage and release, and acting as signaling molecules that regulate cellular processes. Therefore, we are interested in understanding whether fasting reversibly remodels fatty acid chain composition in glycerophospholipids in brain tissues.

We first analyzed the proportions of fatty acid chains in glycerophospholipids from the 4 tissues in *Ad Libitum* chow-fed mice (Fig. 5A and Suppl Fig. 8A). The fatty acid chain compositions in all functional groups are provided in Suppl Fig. 9. It is known that arachidonic acid (AA, 20:4) and docosahexaenoic acid (DHA, 22:6) are the most abundant polyunsaturated fatty acids (PUFAs) in the brain^24^. Consistent with this knowledge, our data demonstrated that AA and DHA make up the biggest proportions of PUFAs (as opposed to saturated FA’s: FA:0 or FA:1) in both HY and BS (Fig. 5A and Suppl Fig. 10A). Surprisingly, the HY contains a much higher proportion of saturated fatty acids compared to the BS, which is driven by the high proportion (almost double; 19.6% vs 36.7%) of palmitic acid (PA, 16:0) in the HY (Fig. 5A and Suppl Fig. 10A). We further analyzed the lipid classes that drove this difference, and identified that it was due to the high abundance of 16:0 fatty acid-containing PC and PC plasmalogen (PC P) in the HY compared to the BS (Suppl Fig. 10B). This is consistent with our previous finding that the HY contained significantly more abundant PC and PC P than the BS (Fig. 2B).

**Fig 5:**
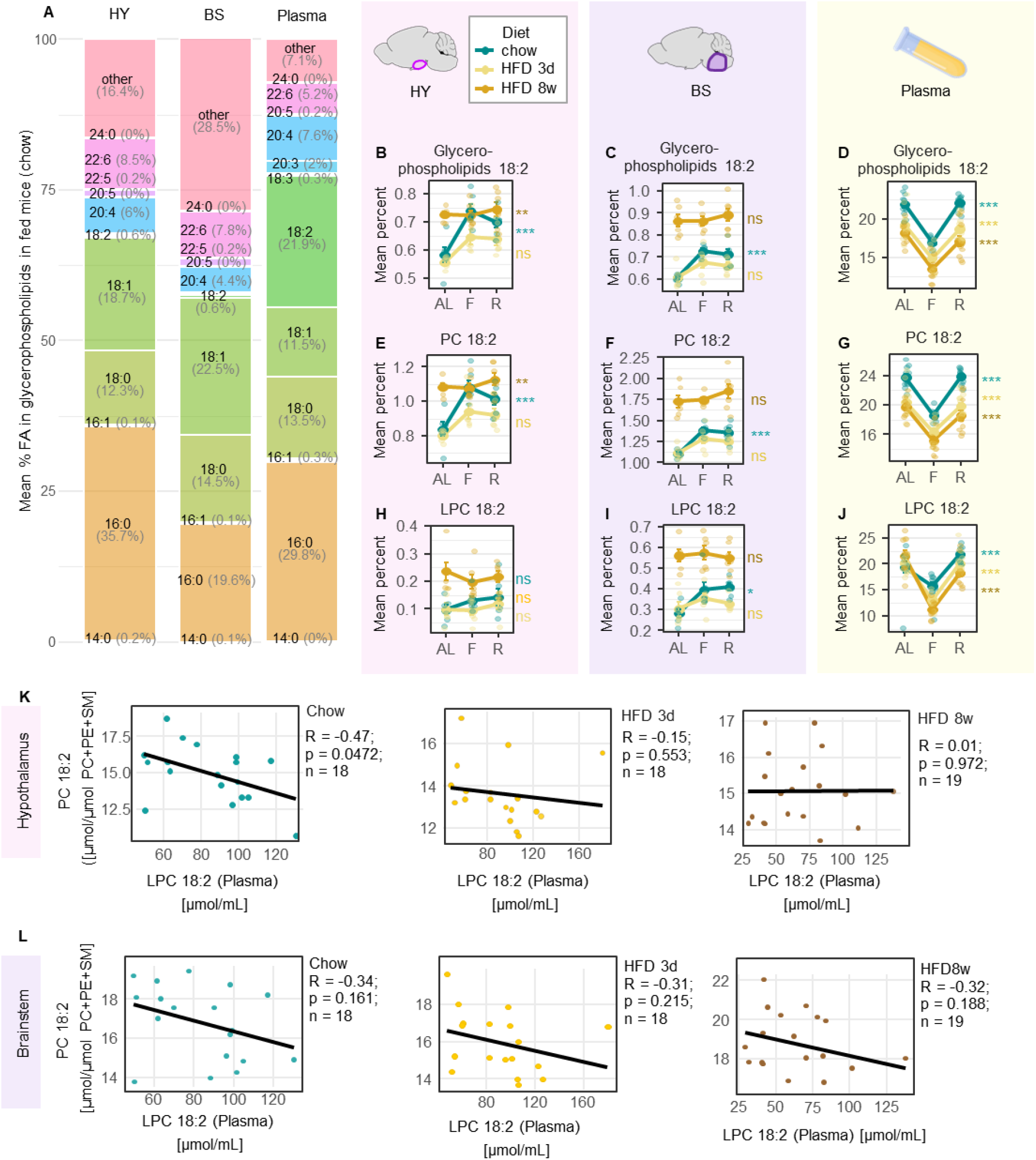
Phospholipids containing fatty acid 18:2 lose elasticity with high fat diet feeding in the hypothalamus. **A** Proportion of lipids containing different fatty acids in hypothalamus (HY), brainstem (BS) and plasma. **B-J** Diet-dependent changes in the relative abundance of lipids containing FA 18:2 across feeding states in HY (left), BS (center) and plasma (right). Top row: percentage of Glycerophospholipids containing 18:2; middle row: Percent PC containing 18:2; bottom row: Percent LPC containing 18:2 **K-L** Correlations of the concentrations of PC 18:2 in hypothalamus (E) and brainstem (F) with LPC 18: 2in plasma in chow (left, cyan), HFD 3d (middle, yellow) and HFD 8w (right, brown). Feeding status (AL = ad libitum fed, F = fasted, R = refed). Diets : chow (cyan), HFD 3d (yellow) and HFD 8w (orange). Statistics: data are represented as mean ± SEM; individual mouse values are overlaid. N= 6-7 mice / feeding status/ diet. One way ANOVA comparing status within each diet; p=*0.05’ **0.01 ***0.001 .

Next, we sought to determine whether fasting reversibly remodels the proportion of any particular fatty acid chains in glycerophospholipids. Our analysis demonstrated that the most dramatic and consistent reversible change with fasting in the HY and BS was the significantly increased proportions of glycerophospholipids containing linoleic acid (LA, 18:2) with fasting, which was reversed by refeeding (Fig. 5B&C). Linoleic acid only represented a small proportion (∼0.6%) of fatty acid chains in glycerophospholipids in both HY and BS, and it was much more abundant in plasma (∼21.9%, Fig. 5A) and CSF (∼12.5%, Suppl Fig. 8A). Interestingly, we observed an opposite and significant change in LA-containing glycerophospholipids with fasting in plasma (Fig. 5D), but not CSF (Suppl Fig. 8B).

To understand which lipid classes drove this global increase of LA-containing glycerophospholipids with fasting in the HY and BS, we analyzed the changes of LA within each lipid class. Our data demonstrated that in both HY and BS, it was the LA-containing PCs that drove the fasting-induced increase of LA-containing glycerophospholipids (Fig. 5E&F). We performed the same analysis in 3d- and 8wk-HFD mice, and found that this fasting-induced increase in LA-containing lipids in the brain was abolished by HFD interventions (Fig. 5B, C, E, F), suggesting the existence of a specific and tunable mechanism that regulates the abundance of these lipids in the brain.

### Linoleic acid (LA)-containing PC abundances in the hypothalamus are strongly correlated with lysophosphatidylcholine 18:2 (LPC 18:2) abundance in plasma

Linoleic acid is an essential PUFA, meaning it must be obtained from the diet, and is a precursor to other PUFAs such as arachidonic acid (AA, 20:4). LA and its derivatives are essential for fetal and neonatal brain development. Since LA cannot be synthetized in mammals, the fasting-induced increase in LA-containing lipids in the brain could be either due to a reduced conversion of LA to its downstream PUFAs, or to an increased uptake of LA from peripheral circulation.

In mammals, LA is desaturated by fatty acid desaturase 1 and 2 (FADS1/FADS2), and elongated by ELOVL fatty acid elongase 2 and 5 (ELOVL2/ELOVL5), to form the downstream PUFAs^25^ (Suppl Fig. 11A). Of note, we did observe reversible decreased expression of the mRNA encoding for these enzymes with fasting in the HY and BS in chow-fed mice (Suppl Fig. 11B), suggesting that fasting might reduce the desaturation and elongation of LA to its downstream PUFAs, and consequently cause an accumulation of LA-containing lipids in the HY. However, animal studies have demonstrated that most PUFA-containing lipids in the brain must be obtained from the circulation, as the synthesis of PUFA-containing lipids in the brain is negligible^26,27^. Moreover, human metabolic studies using stable isotope tracing also showed that the conversion of LA to its downstream PUFAs is complicated and limited, which is typically below 5% in adults^28^. Therefore, we searched for evidence of increased uptake of LA-containing lipids from the circulation.

There are at least 2 pools by which plasma fatty acids supply the brain: free (non-esterified), and esterified in LPCs. It has been shown that fatty acids bound to LPCs, not free fatty acids, are the main form of fatty acids that are transported across the blood-brain barrier (BBB) into the brain, via a transporter known as “major facilitator superfamily domain-containing 2A” (MFSD2A)^29,30^. This is supported by the evidence that dietary LPC 22:6 (LPC-DHA) increased brain DHA-containing lipids, and improved memory in mice, whereas free DHA did not^31^. Interestingly, humans carrying genetic mutations in *MFSD2A* develop lethal microcephaly syndrome associated with inadequate uptake of LPC from the circulation, demonstrated by the fact that those individuals had ∼80% higher total plasma LPC concentrations including LPC 18:2 compared to healthy age-matched controls^30^. After being transported into the brain, the LPCs are remodeled into PCs via LPC acyltransferase 3 (LPCAT3), and those PCs can be converted back to LPCs via phospholipase A2 (PLA_2_), through the Land’s cycle^28^ (Suppl Fig. 11C). To identify whether the reversible increase of LA-containing lipids in the brain induced by fasting is due to an increased uptake of LA from peripheral circulation, we first performed qPCR in the HY and BS to look at the expression of key genes in this pathway. We observed a reversible fasting-induced increase in *Mfsd2a* mRNA expression, and a reversible fasting-induced decrease in *Pla2g4a* mRNA expression in chow-fed mice (Suppl Fig. 11D). These reversible fasting-induced changes were abolished by feeding mice with HFD. We further analyzed LPC 18:2 in plasma, and found out it was decreased by fasting, and this decrease was reversed by refeeding (Fig. 5J). We next performed Pearson correlation analysis, and identified a significant negative correlation (r = -0.47, p = 0.0472) between LA-containing PCs in the HY and LPC 18:2 in the plasma of mice fed on chow (Fig. 5K). Importantly, this correlation was progressively abolished by feeding mice with HFD for 3 days and 8 weeks (r = -0.15 and r = 0.01 respectively) (Fig. 5K). This is consistent with our previous finding that the fasting-induced increase in LA-containing PCs in the HY was abolished by HFD intervention (Fig. 5E). Of note, although similar fasting-induced patterns of LA-containing lipids (Fig. 5C&F) and gene expressions (Suppl Fig. 11D) were observed in the BS, the correlation between LA-containing PCs in the BS and LPC 18:2 in the plasma was not significant in chow-fed mice, and not affected by HFD interventions (Fig. 5L).

The above results suggest that fasting might facilitate the transport of LPC 18:2 from plasma into the HY, via LPC transporter MFSD2A. These LPC 18:2 may then be converted into LA-containing PCs in the HY, which is mediated by the fasting-inhibited expression of PLA_2_. However, this pathway in the BS is not as prominent as the HY, suggesting that it might play a more important role in the HY than the BS in regulating energy balance during fasting.

Taken together, our datasets and extensive analysis of the lipidomes of these tissues identify a significant proportion of lipids in the hypothalamus and brainstem exhibit dynamic remodeling to the metabolic challenge imposed by an AL-F-R cycle, and that this remodeling is diet-dependent. Importantly, fatty acid chain composition analysis identified the reversible fasting-induced increase in linoleic acid (18:2)-containing phosphatidylcholines in both the hypothalamus and brainstem, suggesting a previously unrecognized role for this lipid in the physiological response to fasting.

## Discussion

In this study, we sought to determine whether the lipidome in the hypothalamus and brainstem responds to fasting-induced energy deprivation and returns to its baseline homeostatic state after energy balance is restored. Such “elasticity” would suggest a specific and regulated control of the abundance of particular lipids, implicating them in important biological processes in these brain regions. Our findings showed that 45% and 36% of lipid species in the hypothalamus and brainstem respectively, demonstrated elastic behavior during an AL-F-R cycle. During fasting-induced energy deprivation, plasma lipidome, mostly glycerolipids and fatty acids, showed robust adaptations to serve as energy sources, demonstrated by marked increases in FFAs and acylcarnitine, as well as significant decreases of diacylglycerides (DGs) and TGs. In contrast, only a low proportion of glycerolipids and fatty acids in the hypothalamus and brainstem showed reversible response to fasting-induced energy deprivation. Instead, the majority of elastic lipid species are glycerophospholipids that form cell membranes, regulate membrane permeability and act as cell signaling molecules, suggesting the response of lipidome to an AL-F-R cycle in the hypothalamus and brainstem are more likely to be involved in the molecular control pathways of energy balance, rather than directly serving as energy sources.

High-fat diet (HFD) is a primary driver of obesity and metabolic disorders like insulin resistance, type 2 diabetes, and cardiovascular disease^32^. HFD leads to dysregulation in both the hypothalamus and brainstem, including neuroinflammation as well as insulin and leptin resistance, which often appear before the onset of significant body weight gain^33^. To understand whether the metabolic elasticity of the lipidome in the hypothalamus and brainstem has a potential role in the HFD-induced dysregulation, we compared the LElaS scores of the tissue lipidome from mice that were challenged with short-or long-term HFD interventions, and observed a change from an A-to a V-shape in the elasticity of glycerophospholipids. These elevated proportions of elastic glycerophospholipids in the hypothalamus and brainstem might be involved in membrane remodeling or act as signaling precursors during fasting when mice are fed on a chow diet; however, our data suggested this fasting-induced adaptation might be abolished by long-term high-fat diet feeding.

We observed significant and reversible increase of LA (18:2)-containing PCs induced by fasting in the hypothalamus and brainstem. Interestingly, a similar effect was observed previously in the skeletal muscle of fasted and insulin-deficient mice^34^, where a decrease of DHA-containing PCs was concomitantly observed in skeletal muscle^34^, which we didn’t observe in the brain in our study. As we also observed a significantly decreased LPC 18:2 in plasma upon fasting, as well as a strong negative correlation between hypothalamic LA-containing PCs and plasma LPC 18:2, we propose a potential pathway by which the fasting-induced increase of LA-containing PCs in the hypothalamus might be due to facilitated transport of LPC 18:2 from plasma into the hypothalamus, via LPC transporter MFSD2A. Those LPC 18:2 would then be available to be converted into LA-containing PCs in the hypothalamus. Plasma LPC 18:2 has been reported to be reduced with age^35^, as well as negatively correlated with multiple metabolic disorders such as obesity^36^, type 2 diabetes^36-40^, coronary heart disease^41,42^ and myocardial infarction^43^. Moreover, a recent study in older adults indicated that plasma LPC 18:2 was the only independent predictor of gait speed^44^, which is an important indicator of disability and mortality in older adults. Taken together, plasma LPC 18:2 might play important roles in regulating energy homeostasis and metabolism, via mediating LA-containing PC pathways in the brain and skeletal muscle.

Both the hypothalamus and brainstem contain diverse neuronal and glial cell types organized into distinct nuclei. Importantly, lipid metabolism is handled differentially^45^ and synergistically by these cell populations^46^. Recent studies also revealed distinct lipid profiles among different cell types in the brain, like neurons, microglia, astrocytes and oligodendrocytes^45,47^. Our lipidomics data were collected from homogenates of whole mouse hypothalamus and brainstem, and therefore we were unable to provide information about cell type-specific lipidome elasticity. However, the above-mentioned studies with lipidome data in specific cell types are either based on isolated cells from newborn mice^45^, or iPS (induced pluripotent stem) cell-derived neurons, astrocytes and microglia^47^, making it impossible to perform fasting or diet interventions under these conditions. Moreover, isolating brain cells from adult mice is still challenging, as current protocol of brain cell isolation is based on neonatal mice^48^. In future studies, it is important to understand the lipidome elasticity in specific cell types from different brain regions, which will provide in-depth knowledge in brain lipid metabolism.

Aside from the specific identification of this lipid axis in the brain, our experimental approach has demonstrated that the study of biological elasticity is a novel way to study lipid dynamics. It can capture the remodeling of brain lipidome following metabolic challenges - rather than comparing snapshots of the lipidome under a single metabolic state. Moreover, the elasticity scoring analysis is a unique way to identify lipids with important biological functions that do not differ in their baseline expression between diet interventions.

In summary, this study revealed that around 40% lipids in the hypothalamus and brainstem displayed reservable reversible response to an AL-F-R cycle, demonstrating dynamic remodeling of the brain lipidome in response to metabolic challenges. After long term high fat diet intervention, the pattern of this remodeling was shifted. Fatty acid chain composition analysis revealed a significant and reversible increase in linoleic acid-containing PCs induced by fasting in both the hypothalamus and brainstem. This response was also abolished by high fat diet intervention. Future research is necessary to understand the roles of lipidome remodeling in the regulation of energy metabolism.

## Methods

### Animals

All animal experiments were approved by the A+ Research Alliance (ARA) Animal Ethics Committee (E/2000/2020/B) and conformed to research guidelines of the National Health and Medical Research Council of Australia. Male C57BL/6J mice were either purchased from the Animal Resources Centre in Perth, Australia or bred at the ARA Precinct Animal Centre. Mice were housed at the ARA Precinct Animal Centre at 22^°^C on a 12hr light/dark cycle and had access to a rodent chow (11% energy from fat, Specialty feeds, Australia) and water *ad libitum*, with cages changed weekly. At study end, mice were anesthetized with a lethal dose of ketamine/xylazine prior to blood and tissues collection which were frozen at -80°C for subsequent analysis.

### Diet intervention and fasting-refeeding experiments

Mice were allocated to be either fed chow (11% energy from fat, Specialty Feeds SF00-100, Australia) or high fat diet (45% energy from fat, Specialty Feeds SF04-001, Australia) for 3 days or 8 weeks before being killed at 13 weeks of age (WOA). Before being humanely killed, mice were randomly allocated to be fed *ad libitum*, fasted for 24 hours, or fasted for 24 hours followed by a two-hour refeeding.

### Lipid extraction and sample preparation

Tissues collected include whole hypothalamus (labelled “HY” n=6 per group), brainstem (Pons and medulla, labelled “BS”, n=6 per group), plasma (n=12 per group) and cerebrospinal fluid (labelled “CSF”, n=5-7 per group). Brain tissues (HY and BS) were homogenized and sonicated in PBS with 0.1M DTPA (diethylenetriaminepentaacetic acid) and 0.1M BHT (butylated hydroxytoluene). Approximately 50 µg of protein in 10 µl of solution was transferred to a fresh tube for extraction. 10 µl of plasma were used for extraction. All CSF collected was used for extraction, with volume ranging from 5 to 15 µl, measured upon collection with a Hamilton syringe. Lipid extraction was performed as described previously^49^. In brief, samples were randomized and were mixed with 100 µl of butanol:methanol (1:1) with 10mM ammonium formate which contained a mixture of internal standards. Samples were vortexed thoroughly and set in a sonicator bath for 1 hour maintained at room temperature. Samples were then centrifuged (14,000 g, 10min, 20°C) before transferring into samples vials with glass inserts for analysis.

### Liquid Chromatography Mass Spectrometry

Lipid analysis was performed by liquid chromatography-electrospray ionisation-tandem mass spectrometry (LC-ESI-MS/MS) on an Agilent 1290 Infinity II HPLC coupled to an Agilent 6495C triple quadrupole mass spectrometer updated from our previously described method^50^.

Analysis of lipid extracts were performed on an Agilent 6495C QQQ mass spectrometer with an Agilent 1290 series HPLC system and a single ZORBAX eclipse plus C18 column (2.1×100mm, 1.8µm, Agilent) with the thermostat set at 45°C. Scheduled multiple reaction monitoring was done primarily in positive ion mode. The running solvent consisted of solvents A and B containing of water: acetonitrile: isopropanol in the ratios of 5:3:2 and 1:9:90 respectively, including 10 mM ammonium formate (in solvents A and B) and 5μM medronic acid (in solvent A only). The chromatography settings are as follows: Starting at 15% B, going to 50% at 2.5 minutes, 57% at 2.6 minutes, 70% at 9 minutes, 93% at 9.1 minutes, 96% at 11 minutes, 100% at 11.1 minutes and holding until 12 minutes before going down to 15% at 12.2 minutes and re-equilibrating at 15% B until 16 minutes. The mass spectrometer settings are as follows: gas temperature, 200°c, gas flow, 17L/min, nebuliser pressure, 20psi, sheath gas temperature, 250°c, sheath gas flow, 10L/min, capillary voltage, 3500V, nozzle voltage, 1000V. Positive and low-pressure RF was set to 210 and 110 respectively.

### Quality control samples

The National Institute of Standards and Technology (NIST) human plasma standard reference material 1950 were run as plasma quality control (PQC) samples as previously described^50^. In addition, for brain tissues and CSF, we used identical lipid extracts, which were prepared by pooling the lipid extracts from multiple brain or CSF samples and using this mixture to prepare multiple aliquots which were referred to as brain or CSF quality control (brain QC or CSF QC) samples. For plasma, a PQC was included in every 10 plasma samples. For CSF, a CSF QC was included in every 10 CSF samples. For brain tissues, a PQC and a brain QC were included in every 9 brain samples.

### Data normalization

Quantification of lipids from MS analysis was performed using Mass Hunter Quant B10.0, where lipid abundances were calculated by relating each area under the chromatogram for each lipid species to the corresponding internal standard (ITSD). Blanks with internal standards were used to determine background levels of each lipid, and were used for subtraction.

### Data inclusion and exclusion criteria

Lipids with more than 50% of samples with NAs for any tissue-diet combination were removed, and the rest of the data was imputed with half the minimum value for any given lipid. To detect potential technical outliers in the datasets, we first assessed each dataset using multiple unbiased approaches. For each tissue, we generated heatmaps of log-transformed lipid proportions and applied hierarchical clustering. Heatmaps were then visually screened to flag any samples with atypical profiles. In parallel, we performed principal component analysis (PCA) for each group and manually reviewed the score plots to support outlier detection. Individual lipid concentrations higher or lower than 4IQ were qualified as ultra outliers and were deleted from the dataset.

### Lipid nomenclature

Lipid nomenclature was assigned according to the LIPID MAPS consortium^51-53^. Phospholipids with resolved structural detail characteristics (i.e.known acyl-chain composition) are reported as ‘PC(16:0_20:4)’, with PC being the lipid class and (16:0_20:4) indicating the acyl chains attached to the glycerol backbone, irrespective of sn1/ sn2 position. When specific structural annotations are not available, lipids are named based on their sum acyl chain length and degrees of saturation (e.g., PC(36:4)). For fatty acid chain annotation, lipids without specific annotations were labeled as “other”, and their concentration was multiplicated by the number of fatty acid chains present in their structure to account for these lipids in the percent fatty acid abundance. Isomeric lipid species separated chromatographically but incompletely annotated were designated ‘a’, ‘b’ and so on, with ‘a’ and ‘b’ representing the elution order. Owing to technical limitations, we were unable to assign acyl chains to a specific sn1/ sn2 position for the majority of phospholipid species; only that the acyl chain was present at either the sn1 or sn2 position. In the case of either PC and PE, the alkyl or alkenyl chains are always located at the sn1 position and the acyl chain is always at the sn2 position. Fatty acid chains in TGs were quantified by neutral-loss analysis.

### Data analysis

Statistical analysis of lipidomic data was conducted using R (4.2.2) as previously described^50^. The lipidomics data was normalized to total membrane lipids (sum of PC, PE and SM) and reported as pmol per µmol of membrane lipids. and Log10 transformed prior to statistical analysis unless stated otherwise. Z-scores were calculated by using the R built-in scale() function.

One-way ANOVAs were conducted in R using built-in functions, followed by Tukey’s post-hoc tests. ANOVA p-values were adjusted for multiple comparisons using the Benjamini–Hochberg procedure to control the false discovery rate^54^. FDR-corrected P values were statistically significant when p< 0.05.

To perform PCA analysis (Fig. 1), we identified lipids consistently detected across all tissue–diet groups and retained only these shared species. The filtered datasets were merged, the remaining missing values were imputed using a half-minimum approach, and the data were log10-transformed prior to analysis. Principal component analysis (PCA) was subsequently performed on the resulting matrix of 511 lipid species, using the PCA() function from the FactoMineR package.

Total lipid class abundances were derived from the summation of all lipid species within each class. In Suppl Fig. 1, classes were grouped into functional groups, and the percent abundance of each class within each functional group/ tissue combination was calculated. Low abundance classes were labelled as “low” and plotted separately as a percent proportion of low abundance classes. In Fig. 2, totals were normalized to volume (µmol/mL), and brain tissue concentrations were normalized to PC+PE+SM and z-scored.

The Elasticity scoring system was designed to measure an organism’s capacity to cope with and recover from metabolic fluctuations^21^. The Lipid Elastic Scores (LElaS) were adapted from Gene Elasticity scores^21^. Briefly, the changes in lipid species concentrations from ad lib (AL) to fasted (F) and F to refed (R) within each diet condition were compared using ANOVA followed by post-hoc multiple comparisons, and the resulting fold changes and p-values were used to compute elasticity scores for individual lipid species. As opposed to the original scoring system, here, lipids decreasing with fasting were assigned a negative elasticity score. Lipids were consequently assigned a “shape category” based on their profile in a AL-F-R cycle: Lipid species steadily increasing or decreasing with fasting and refeeding were assigned, “up” or “down”, respectively; lipid species increasing with fasting and decreasing with refeeding were assigned an “A” shape, and vice-versa, a “V” shape; and lastly, lipid species with an absolute elasticity score lower than 0.01 were categorized as “not changed” (NC).

Chi-square and Fisher’s exact tests were used to assess differences in the distribution of categorical lipid counts between experimental group (figure 4, supplemental figures 3 and 7) using base R functions chisq.test() and fisher.test(). For each analysis, lipid species were aggregated into contingency tables (e.g., group × category), and an overall chi-square test of independence was applied when expected cell counts were sufficiently large (generally ≥5 in most cells). When one or more expected counts fell below this threshold, Fisher’s exact test was used instead to provide an exact p-value. Where post-hoc pairwise comparisons were performed, the same decision rule (chi-square vs Fisher’s exact) was applied to each 2×2 table, and p-values were corrected for multiple testing using the Benjamini–Hochberg procedure.

Fatty acid chain analysis (Fig. 5, Suppl Fig. 7-10): For each tissue, lipid species present in the three diets were filtered. Fatty acid composition was quantified within each tissue in different diets and feeding status by aggregating fatty acyl–resolved lipid species within each lipid functional group (or class) and fatty acid type. Fatty acid types were defined as follows: “regular fatty acids” (labelled “FA”, 14:0 to 24:1), P- and O-linked fatty acids (labelled, respectively, P-FA and O-FA), methylated fatty acids (FA;Me), and Oxidized fatty acids (FA;O). For each tissue × diet × feeding status × functional group (or class) × fatty acid type combination, the concentration of the lipid species containing individual fatty acid were summed and expressed as the percentage contribution of each fatty acid relative to the total fatty acid signal within that FA type (i.e., each fatty acid type was normalized to 100%). For lipid species for which individual fatty acyl chains could not be resolved; the measured lipid concentration was multiplied by the annotated number of fatty acids and the resulting total was assigned to “other”. Cardiolipins (CL) were excluded from the glycerophospholipid subset because their acyl-chain composition could not be completely resolved with our annotation approach. The resulting percent abundances for each fatty acid were compared between feeding statuses using one way ANOVA individually for each diet.

Pearson correlation in Fig. 5 was performed on concentrations of brain PC 18:2 ([μmol/μmol PC+PE+SM]) and plasma LPC18:2 [μmol/mL] using base R cor() function.

### qPCR

Quantitative real-time PCR (qPCR) was performed on a Quant Studio 7 Real-Time PCR system (Applied Biosystems) using the iTaq Universal Sybr Green Kit (Bio-Rad). Relative gene expression levels were calculated using the ΔΔCt method with *Csnk2n2* as the reference gene. The primer sequences are provided in Table 1. Expression was normalized to gene expression of mice on chow (fed status).

**TABLE 1:**
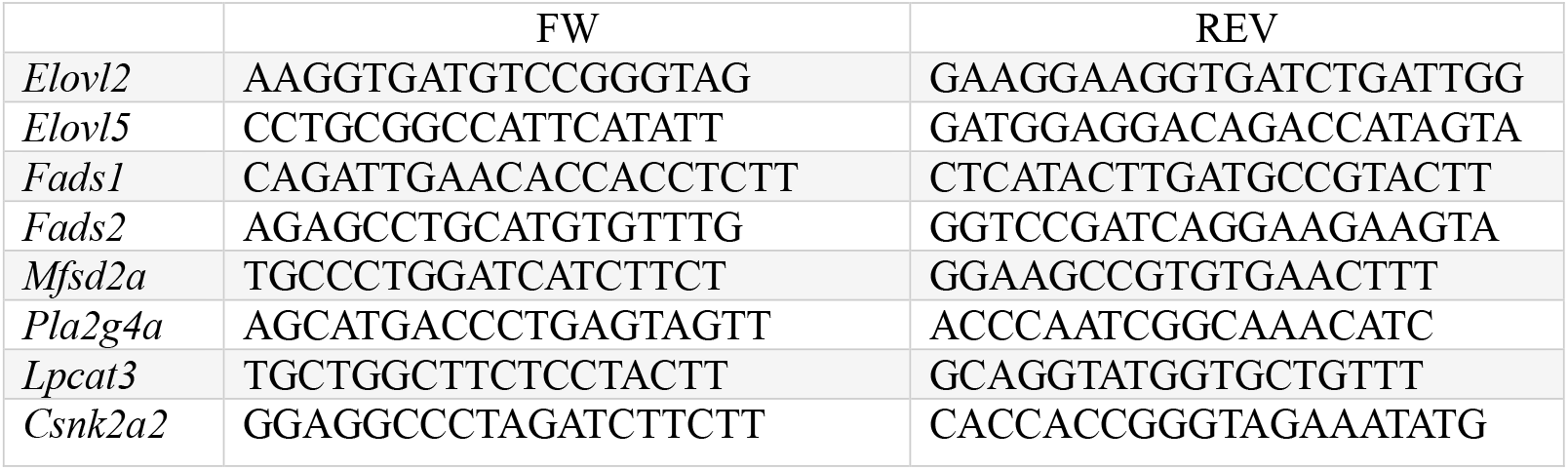
Primer sequences.

## Data availability

Data can be requested to the Lead contact.

## Author Contributions

YW, AB, KH and BGD conceived and designed the study. AB, JXF, HYW, HL, YL, NM and YW performed experiments. YW, AB, KH and BGD analyzed and interpreted data. PJM provided lipidomics expertise. BGD provided access to resources and experimental advice. YW, AB and BGD wrote the manuscript. All authors read and approved the manuscript.

## Acknowledgements

This work was funded in part by NHMRC Ideas grants (1186512, 2019687). BGD has received funding support from the National Heart Foundation of Australia Future Leader Fellowship (101789) and NHMRC of Australia (Investigator Grant 2016530). KH is supported by an NHMRC Investigator grant (1197190).

We thank all members of the MMA laboratories at Baker Institute for their support.

