## Supplemental figures for "Diet- and metabolic state-dependent remodeling of the mouse brain lipidome"

Adélaïde Bernard[1,2], Kevin Huynh[1,2,3,4], Jen Xin Fach[1,3], Hang Yi Woo[1,3], He Liu[1], Yingying Liu[1], Natalie Mellet[1], Peter Meikle[1,2,3,4], Brian G Drew[1,2,3,4,5], Yi Wang[1,2,3,5]

[1]*Baker Heart and Diabetes Institute, Melbourne, VIC, Australia*

[2]*Baker Department of Cardiometabolic Health, University of Melbourne, Parkville, VIC, Australia*

[3]*School of Translational Medicine, Monash University, Melbourne, VIC, Australia*

[4]*Baker Department of Cardiovascular Research Translation and Implementation, La Trobe University, Melbourne, VIC, Australia*

[5]*These authors contributed equally*

*Author for Correspondence*

**

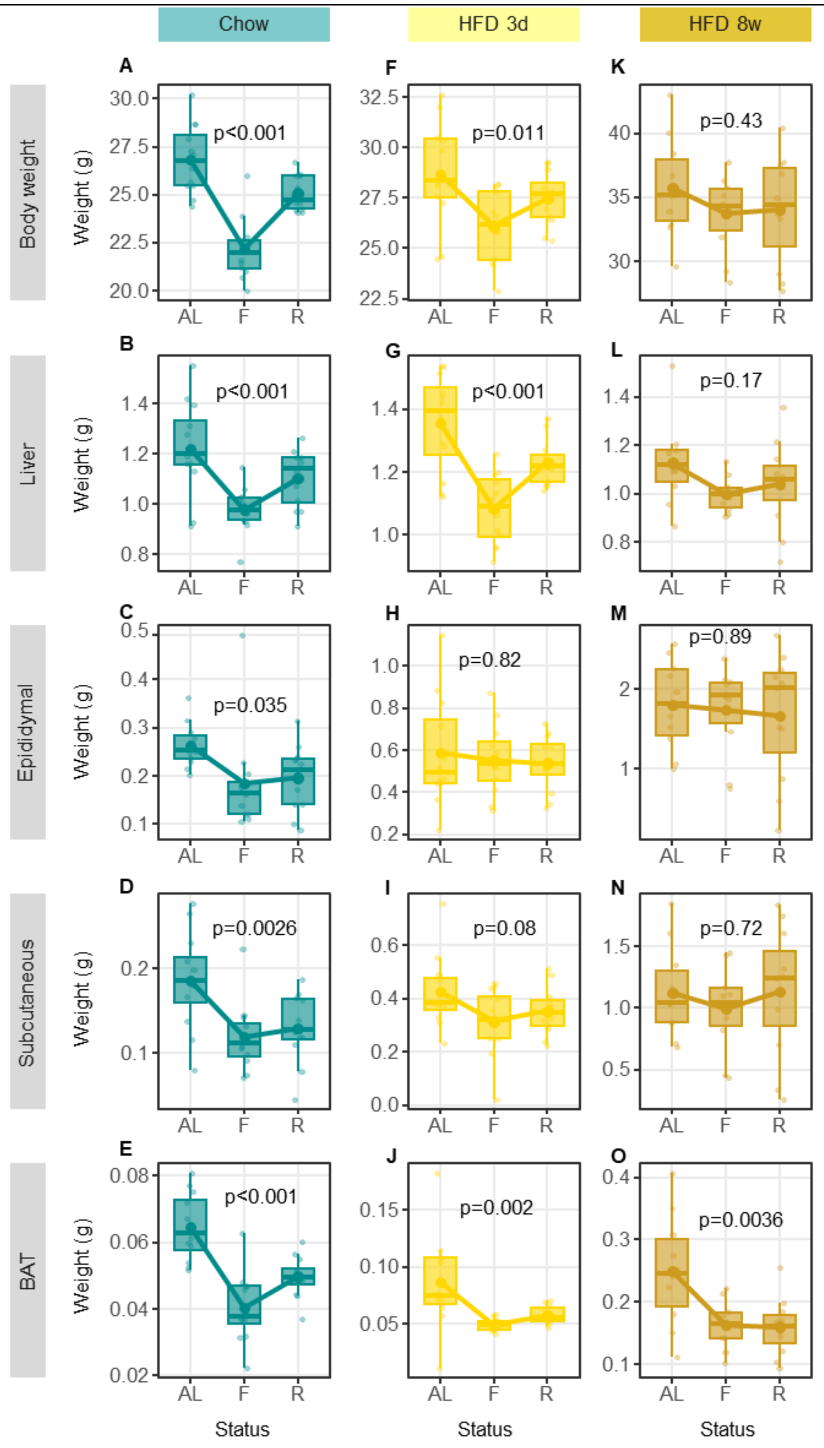


**Supplemental Figure 1. Mouse body and organ weights across metabolic status and diet.** Body weight (**A,F,K**) and organ weights (**B-O**) of all mice involved in the study are represented across feeding status ("AL", ad libitum fed; "F", fasted; "R", refed).) and diets (columns, chow (cyan), HFD 3d (yellow), HFD 8w (orange)). Organs: Liver; epididymal adipose tissue; subcutaneous adipose tissue; and "BAT", brown adipose tissue. Each dot represents an individual mouse. Box-and-whisker plots: boxes show the median (center line) and interquartile range (25th–75th percentiles); whiskers extend to 1.5× the interquartile range. Lines and points overlay the mean for each status group. Sample size, n = 12 mice. One-way ANOVA p-values are displayed above each panel (computed separately within each diet).

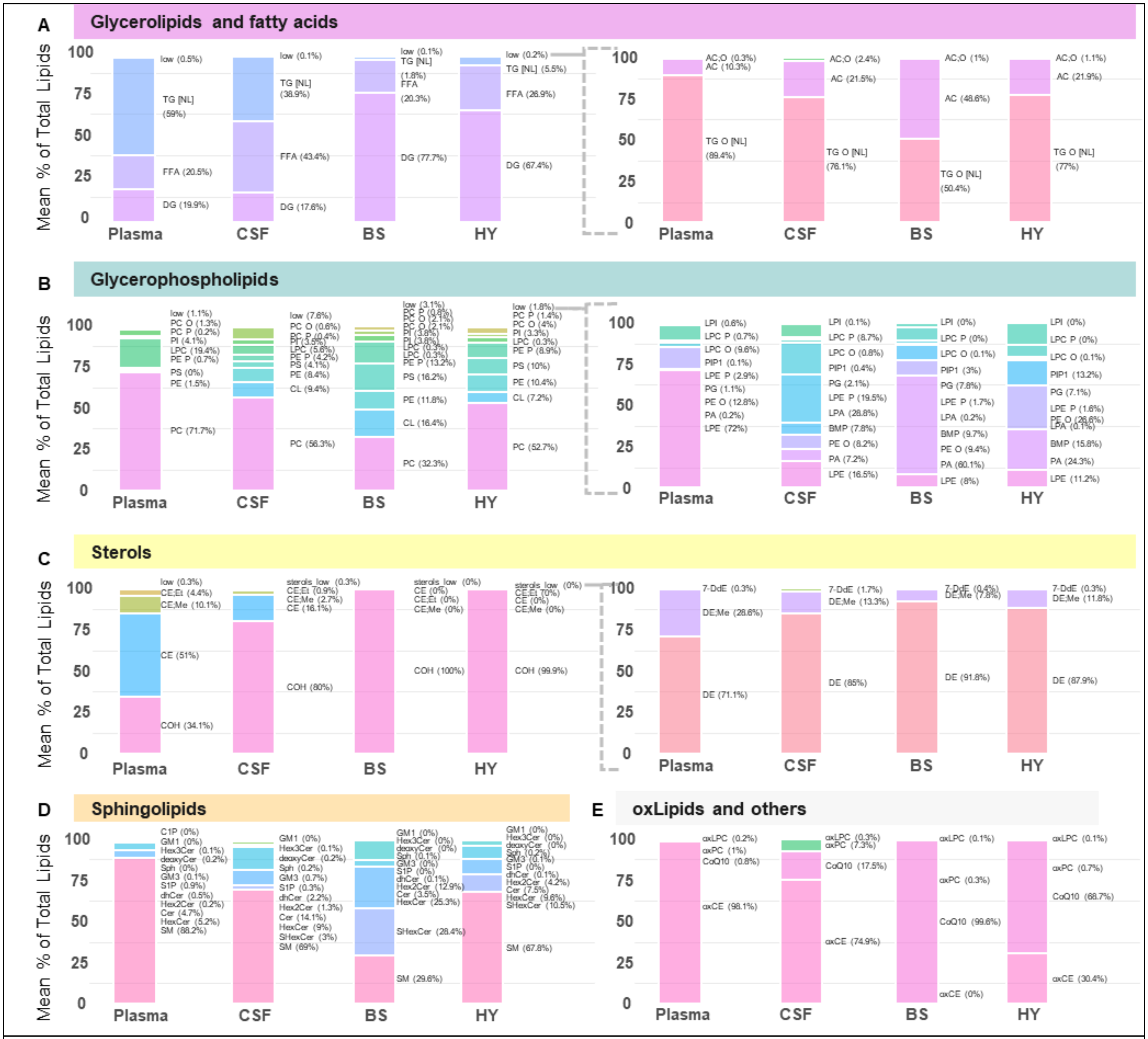


**Supplemental Figure 2: Proportions of lipid classes, split by functional group across tissues.** Percent of total concentrations of measured lipid classes in Plasma, Cerebrospinal fluid (CSF), brainstem (BS) and hypothalamus (HY), organized by functional groups: Glycerolipids and fatty acids (**A**), Glycerophospholipids (**B**), Sterols (**C**), Sphingolipids (**D**) and oxylipids and others (**E**). High abundance lipids are plotted on the left and low abundance lipids on the right (**A-C**).

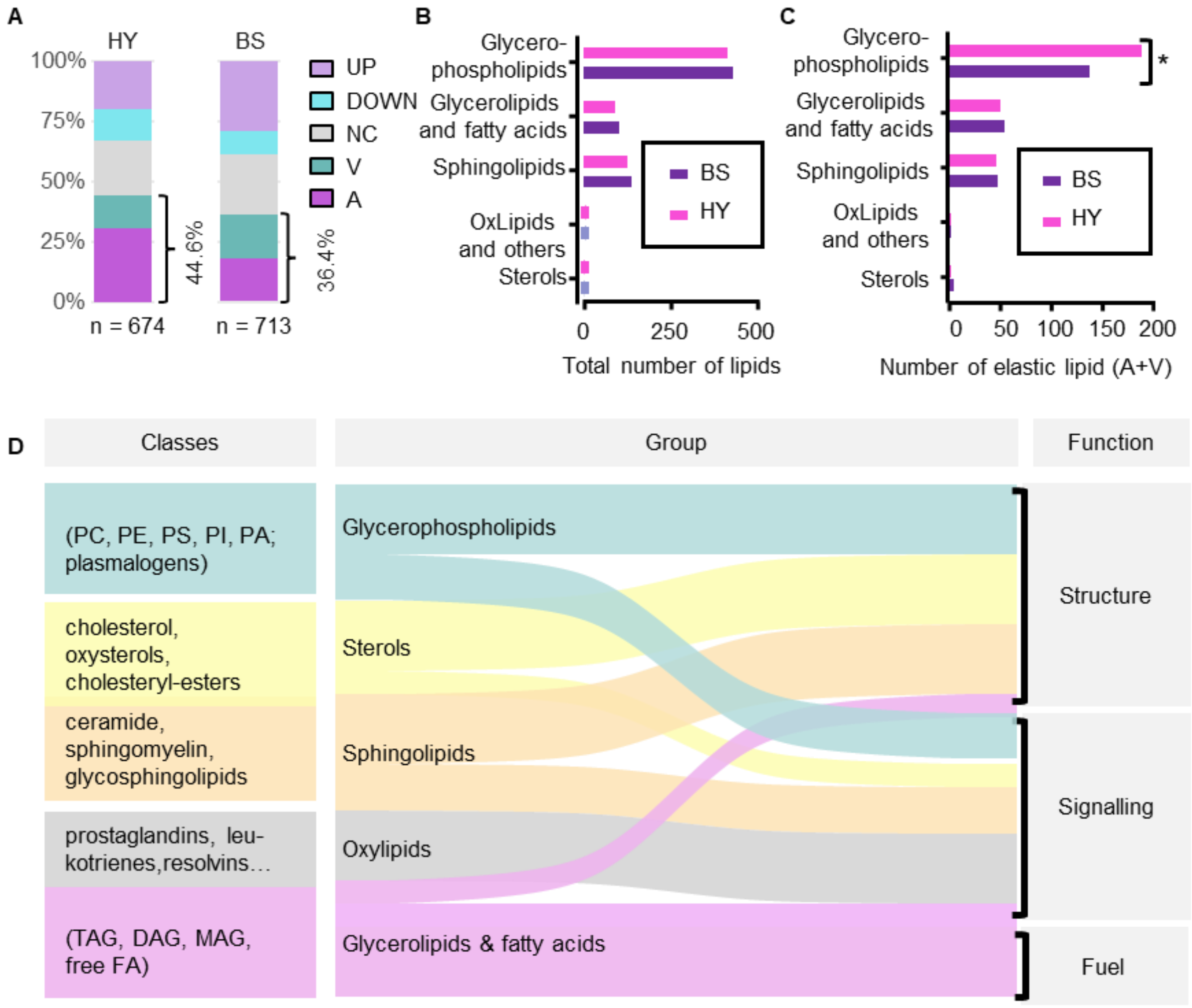


**E**

| Top 20 most elastic lipids | | | |
|---|---|---|---|
| Plasma | CSF | BS | HY |
| CE 18:2;O | LPI 0:0/20:4 | BMP 16:0_18:0 | BMP 18:0_18:1 |
| PC 34:2;O | PC O-44:6 | BMP 18:1_18:2 | BMP 18:2_22:6 |
| Cer 20:1;O2/22:0 | PI 18:1_18:2 | BMP 18:2_22:6 | CL 36:3_36:1 |
| Hex-Cer 18:2;O2/18:0 | CE 16:0 | LPC 0:0/19:0 [a] | CL 36:3_38:4 |
| AC 14:0 | CE 20:0 | LPC 17:1/0:0 [b] | LPA 18:1 |
| AC 15:0 [b] | CE 20:4 | LPC 20:5/0:0 | LPC 0:0/24:0 |
| AC 16:0 | CE 20:5 | LPI 0:0/18:1 | LPC P-17:0/0:0 [a] |
| AC 16:0;O | CE 24:5 | LPI 0:0/18:2 | LPC P-17:0/0:0 [b] |
| AC 16:1 | CE;Me 18:2 \| SE 28:1/18:2 | LPI 18:0 | LPC P-18:1/0:0 |
| AC 16:1;O | SHexCer 18:1;O2/16:0 | LPI 18:2/0:0 | LPI 18:2/0:0 |
| AC 17:0 | AC 22:5;O | PE P-15:0/22:6 [b] | PC 35:5 |
| AC 18:1 | FFA 22:4 | LPC 22:6;O | PC P-16:0/20:5 |
| AC 18:1;O | TG 48:3 [NL-18:3] | CE 22:5 | CE;Me 20:4 \| SE 28:1/20:4 |
| AC 18:2 | TG 50:2 [NL-14:0] | CE;Et 22:6 \| SE 29:1/22:6 | AC 18:0;O |
| AC 18:3 | TG 50:2 [NL-18:2] | Cer 18:2;O2/16:0 | AC 18:1;O |
| AC 20:5 | TG 50:3 [NL-18:2] | Cer 19:1;O2/22:0 | AC 26:1 |
| AC 22:6 | TG 50:4 [NL-20:4] | AC 22:5 | DG 18:1_20:3 |
| TG 50:3 [NL-18:3] | TG 52:5 [NL-18:3] | TG 52:2 [NL-18:2] | FFA 20:2 |
| TG 54:5 [NL-18:3] | TG 56:7 [NL-22:5] | TG 53:2 [NL-17:1] | TG 58:9 [NL-22:6] |
| TG 54:6 [NL-18:3] | TG O-50:1 [NL-17:1] | TG 56:9 [NL-22:6] | TG O-54:3 [NL-17:1] |

**Supplemental Figure 3**: **Elasticity across different lipid functional groups in different tissues.**

**A** Proportion of lipid species belonging to different shape categories in hypothalamus (HY) compared to brainstem (BS).

**B** Total number of lipids from different functional groups in brainstem (BS) and hypothalamus (HY).

**C** Comparison of the number of elastic lipids per functional group in the hypothalamus (HY) compared to brainstem (BS).

**D** Ribbon diagram connecting classes to their corresponding lipid groups and their involvement in different functions (simplified )

**E** Top 20 most elastic lipids in plasma, CSF, brainstem (BS) and hypothalamus (HY), colour-coded by functional group as described in A.

Statistics: Overall differences in lipid-category composition between BS and HY were assessed in A and C by a chi-square test of independence. For each category, enrichment in BS vs HY was evaluated using 2×2 tests (category vs all other categories); Fisher's exact test was used when expected counts were $<5$, otherwise chi-square. P* 0.05

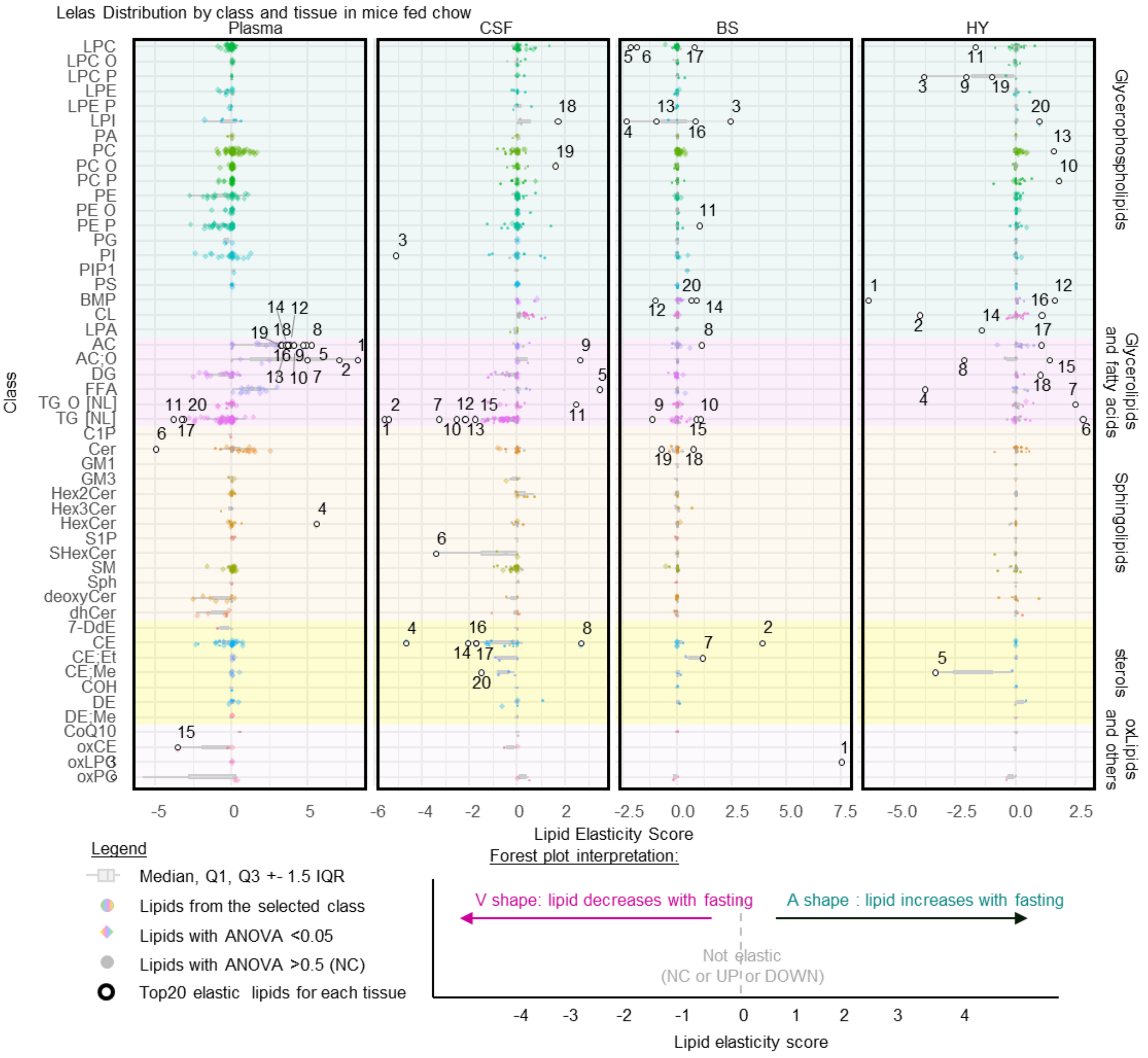


| chow | | | | | | | |
|---|---|---|---|---|---|---|---|
| Plasma | | CSF | | BS | | HY | |
| Ranking | lipid | Ranking | lipid | Ranking | lipid | Ranking | lipid |
| 1 | AC 16:1;O | 1 | TG 48:3 [NL-18:3] | 1 | LPC 22:6;O | 1 | BMP 18:0_18:1 |
| 2 | AC 18:1;O | 2 | TG 56:7 [NL-22:5] | 2 | CE 22:5 | 2 | CL 36:3_36:1 |
| 3 | PC 34:2;O | 3 | PI 18:1_18:2 | 3 | LPI 18:2/0:0 | 3 | LPC P-17:0/0:0 [a] |
| 4 | Hex-Cer 18:2;O2/18:0 | 4 | CE 20:0 | 4 | LPI 0:0/18:1 | 4 | FFA 20:2 |
| 5 | AC 16:1 | 5 | FFA 22:4 | 5 | LPC 0:0/19:0 [a] | 5 | CE;Me 20:4 \| SE 28:1/20:4 |
| 6 | Cer 20:1;O2/22:0 | 6 | SHexCer 18:1;O2/16:0 | 6 | LPC 17:1/0:0 [b] | 6 | TG 58:9 [NL-22:6] |
| 7 | AC 16:0;O | 7 | TG 50:4 [NL-20:4] | 7 | CE;Et 22:6 \| SE 29:1/22:6 | 7 | TG O-54:3 [NL-17:1] |
| 8 | AC 18:1 | 8 | CE 24:5 | 8 | AC 22:5 | 8 | AC 18:0;O |
| 9 | AC 18:2 | 9 | AC 22:5;O | 9 | TG 53:2 [NL-17:1] | 9 | LPC P-18:1/0:0 |
| 10 | AC 18:3 | 10 | TG 52:5 [NL-18:3] | 10 | TG 56:9 [NL-22:6] | 10 | PC P-16:0/20:5 |
| 11 | TG 54:5 [NL-18:3] | 11 | TG O-50:1 [NL-17:1] | 11 | PE P-15:0/22:6 [b] | 11 | LPC 0:0/24:0 |
| 12 | AC 14:0 | 12 | TG 50:2 [NL-14:0] | 12 | BMP 16:0_18:0 | 12 | BMP 18:2_22:6 |
| 13 | AC 22:6 | 13 | TG 50:3 [NL-18:2] | 13 | LPI 18:0 | 13 | PC 35:5 |
| 14 | AC 17:0 | 14 | CE 20:4 | 14 | BMP 18:2_22:6 | 14 | LPA 18:1 |
| 15 | CE 18:2;O | 15 | TG 50:2 [NL-18:2] | 15 | TG 52:2 [NL-18:2] | 15 | AC 18:1;O |
| 16 | AC 20:5 | 16 | CE 20:5 | 16 | LPI 0:0/18:2 | 16 | CL 36:3_38:4 |
| 17 | TG 54:6 [NL-18:3] | 17 | CE 16:0 | 17 | LPC 20:5/0:0 | 17 | AC 26:1 |
| 18 | AC 16:0 | 18 | LPI 0:0/20:4 | 18 | Cer 18:2;O2/16:0 | 18 | DG 18:1_20:3 |
| 19 | AC 15:0 [b] | 19 | PC O-44:6 | 19 | Cer 19:1;O2/22:0 | 19 | LPC P-17:0/0:0 [b] |
| 20 | TG 50:3 [NL-18:3] | 20 | CE;Me 18:2 \| SE 28:1/18:2 | 20 | BMP 18:1_18:2 | 20 | LPI 18:2/0:0 |

**Supplemental figure 4:** Forest plot of Lipid Elasticity scores of all lipids organized by class (Y axis) and tissue (X Axis) in mice fed chow. Top 20 elastic lipids for each tissue are provided at the bottom

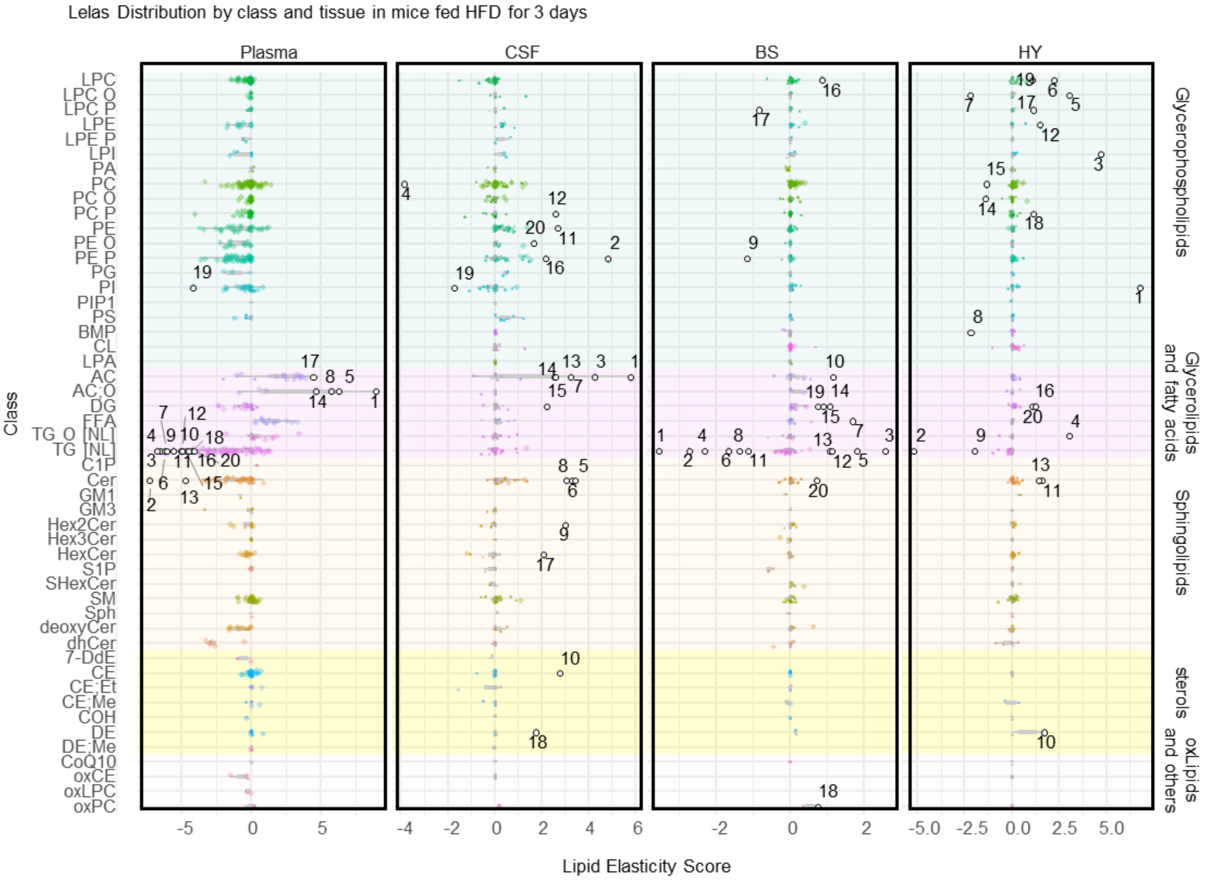


| HFD 3 days | | | | | | | |
|---|---|---|---|---|---|---|---|
| Plasma | | CSF | | BS | | HY | |
| Ranking | lipid | Ranking | lipid | Ranking | lipid | Ranking | lipid |
| 1 | AC 20:3;O | 1 | AC 14:0 | 1 | TG 52:3 [NL-18:2] | 1 | PI 16:0;Me_18:2 \| PI 17:0_18:2 |
| 2 | Cer 16:1;O2/16:0 | 2 | PE P-19:0/20:4 [b] | 2 | TG 54:6 [NL-20:4] | 2 | TG 52:5 [NL-20:4] |
| 3 | TG 52:1 [NL-18:1] | 3 | AC 18:1 | 3 | TG 50:4 [NL-20:4] | 3 | LPI 18:1/0:0 |
| 4 | TG 48:0 [NL-18:0] | 4 | PC 44:4 [b] | 4 | TG 51:2 [NL-17:0] | 4 | TG O-50:1 [NL-15:0] |
| 5 | AC 16:1;O | 5 | Cer 18:2;O2/24:1 | 5 | TG 54:6 [NL-20:5] | 5 | LPC O-20:0/0:0 |
| 6 | TG 50:1 [NL-14:0] | 6 | Cer 18:1;O2/18:0 | 6 | TG 52:2 [NL-18:2] | 6 | LPC 19:0/0:0 [a] (PI-104) |
| 7 | TG 52:1 [NL-18:0] | 7 | AC 16:0 | 7 | FFA 18:3 | 7 | LPC O-24:0/0:0 |
| 8 | AC 18:1;O | 8 | Cer 18:2;O2/18:0 | 8 | TG 54:0 [NL-18:0] | 8 | BMP 16:0_16:1 |
| 9 | TG 54:1 [NL-18:1] | 9 | Hex2Cer 18:1;O2/24:0 | 9 | PE P-15:0/20:4 [a] | 9 | TG 52:2 [NL-18:2] |
| 10 | TG 50:0 [NL-18:0] | 10 | CE 22:4 | 10 | AC 22:6 | 10 | DE 22:6 \| SE 27:2/22:6 |
| 11 | TG 48:1 [NL-18:1] | 11 | PE 17:0_20:4 | 11 | TG 54:3 [NL-18:2] | 11 | Cer 17:1;O2/24:1 |
| 12 | TG 54:0 [NL-18:0] | 12 | PC P-35:2 [a] | 12 | TG 56:9 [NL-22:6] | 12 | LPE 0:0/18:2 |
| 13 | Cer 19:1;O2/16:0 | 13 | AC 20:5 | 13 | TG 56:8 [NL-22:6] | 13 | Cer 20:1;O2/23:0 |
| 14 | AC 16:0;O | 14 | AC 18:0 | 14 | DG 18:2_22:6 | 14 | PC O-42:5 [b] |
| 15 | TG 51:1 [NL-17:0] | 15 | DG 18:0_20:4 | 15 | DG 18:1_20:5 | 15 | PC 39:5 [a] |
| 16 | TG 53:2 [NL-17:1] | 16 | PE P-16:0/20:3 [b] | 16 | LPC 0:0/19:0 [a] | 16 | DG 18:2_22:6 |
| 17 | AC 16:1 | 17 | Hex-Cer 18:2;O2/20:0 | 17 | LPC P-20:0/0:0 | 17 | LPC P-17:0/0:0 [b] |
| 18 | TG 49:1 [NL-17:1] | 18 | DE 18:1 \| SE 27:2/18:1 | 18 | PC 36:4;O | 18 | PC P-16:0/18:3 |
| 19 | PI 18:0_18:1 | 19 | PI 20:0_20:4 | 19 | DG 18:1_22:5 | 19 | LPC 0:0/26:0 |
| 20 | TG 50:1 [NL-18:1] | 20 | PE O-16:0/22:4 | 20 | Cer 20:1;O2/26:0 | 20 | DG 18:2_20:4 |

**Supplemental figure 5:** Forest plot of Lipid Elasticity scores of all lipids organized by class (Y axis) and tissue (X Axis) in mice fed HFD for 3 days. Top 20 elastic lipids for each tissue are provided at the bottom.

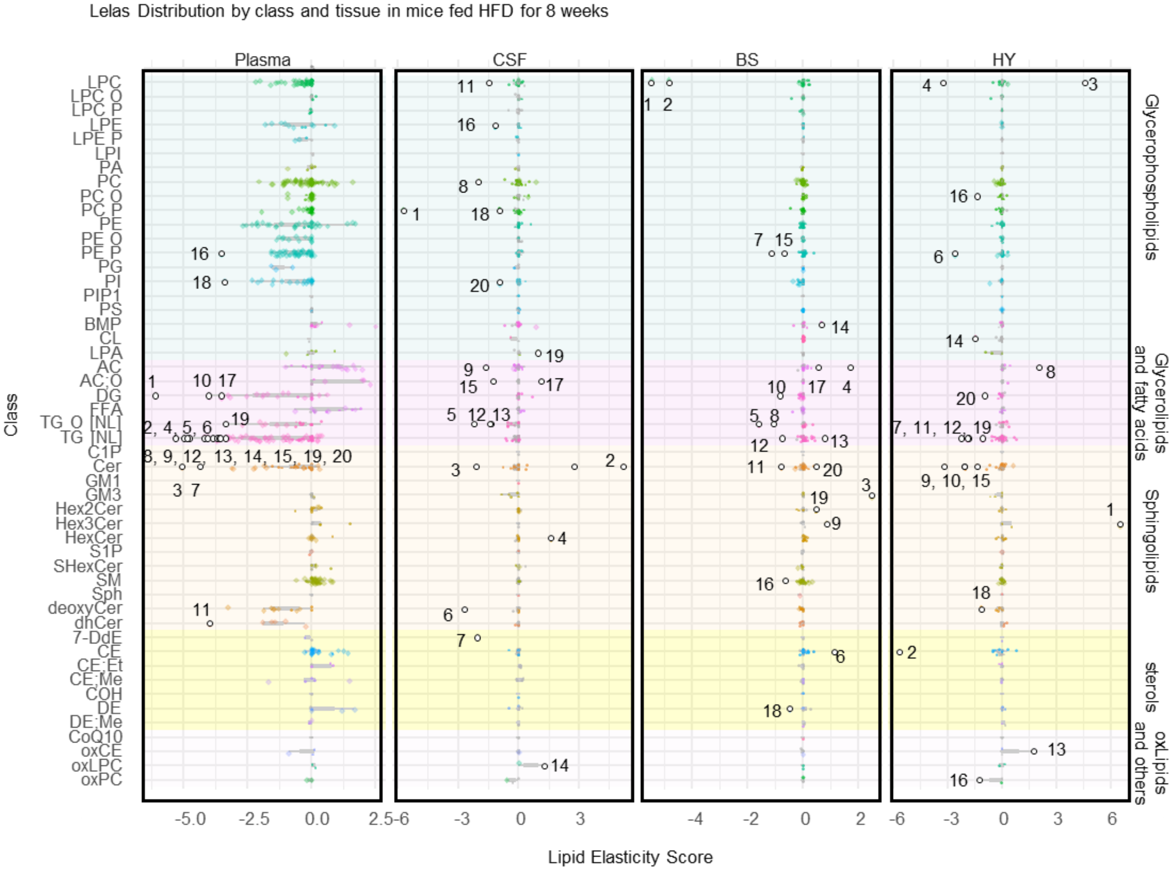


| HFD 8weeks | | | | | | | |
|---|---|---|---|---|---|---|---|
| Plasma | | CSF | | BS | | HY | |
| Ranking | lipid | Ranking | lipid | Ranking | lipid | Ranking | lipid |
| 1 | DG 14:0_16:0 | 1 | PC P-35:2 [a] | 1 | LPC 19:1 [c] | 1 | Hex3Cer 18:1;O2/20:0 |
| 2 | TG 52:1 [NL-18:1] | 2 | Cer 20:1;O2/24:1 | 2 | LPC 0:0/19:0 [a] | 2 | CE 16:0 |
| 3 | Cer 16:1;O2/16:0 | 3 | Cer 17:1;O2/23:0 | 3 | GM3 18:2;O2/24:1 | 3 | LPC 0:0/26:0 |
| 4 | TG 50:1 [NL-14:0] | 4 | Cer 18:0;O/23:0 | 4 | AC 26:0 | 4 | LPC 17:1/0:0 [b] |
| 5 | TG 52:1 [NL-18:0] | 5 | TG O-54:2 [NL-18:1] | 5 | TG O-54:4 [NL-18:2] | 5 | Cer 17:1;O2/22:0 |
| 6 | TG 48:0 [NL-18:0] | 6 | Cer 18:2;O2/23:0 | 6 | CE 18:1 | 6 | PE P-15:0/22:6 [b] |
| 7 | Cer 16:1;O2/23:0 | 7 | 7-DdE 20:4 \| SE 27:3/20:4 | 7 | PE P-15:0/22:6 [a] | 7 | TG 54:6 [NL-18:3] |
| 8 | TG 48:1 [NL-18:1] | 8 | PC 35:5 | 8 | TG O-50:3 [NL-18:2] | 8 | AC 18:2 |
| 9 | TG 54:1 [NL-18:1] | 9 | AC 17:0 | 9 | Hex3Cer 18:1;O2/22:0 | 9 | Cer 16:1;O2/24:0 |
| 10 | DG 16:0_16:0 | 10 | Hex-Cer 16:1;O2/18:0 | 10 | DG 14:0_16:0 | 10 | Cer 19:1;O2/24:0 |
| 11 | Cer 18:0;O2/16:0 | 11 | LPC 17:1/0:0 [b] | 11 | Cer 19:1;O2/20:0 | 11 | TG 51:1 [NL-17:0] |
| 12 | TG 50:0 [NL-18:0] | 12 | TG O-54:4 [NL-18:2] | 12 | TG 54:6 [NL-20:5] | 12 | TG 48:3 [NL-18:3] |
| 13 | TG 51:1 [NL-17:0] | 13 | TG O-52:1 [NL-18:1] | 13 | TG 49:1 [NL-17:1] | 13 | CE 22:6;O |
| 14 | TG 53:2 [NL-17:1] | 14 | LPC 22:6;O | 14 | BMP 16:0_22:6 | 14 | CL 40:7_34:1 |
| 15 | TG 54:2 [NL-18:0] | 15 | AC 24:1;O | 15 | PE P-16:0/18:3 | 15 | Cer 16:1;O2/24:1 |
| 16 | PE P-15:0/20:4 [a] | 16 | LPE 18:2/0:0 | 16 | SM 19:1;O2/24:1 | 16 | PC O-46:7 [a] |
| 17 | DG 16:0_18:1 | 17 | AC 16:1;O | 17 | AC 20:3 [a] | 17 | PC 36:4;O |
| 18 | PI 18:0_18:1 | 18 | PC P-35:2 [b] | 18 | DE 16:0 \| SE 27:2/16:0 | 18 | Cer 18:0;O/24:1 |
| 19 | TG O-54:4 [NL-17:1] | 19 | LPA 20:4 | 19 | Hex2Cer 18:2;O2/16:0 | 19 | TG 52:2 [NL-18:2] |
| 20 | TG 48:2 [NL-18:2] | 20 | PI 36:2 | 20 | Cer 18:2;O2/21:0 | 20 | DG 14:0_16:0 |

**Supplemental figure 6:** Forest plot of Lipid Elasticity scores of all lipids organized by class (Y axis) and tissue (X Axis) in mice fed HFD for 8weeks. Top 20 elastic lipids for each tissue are provided at the bottom

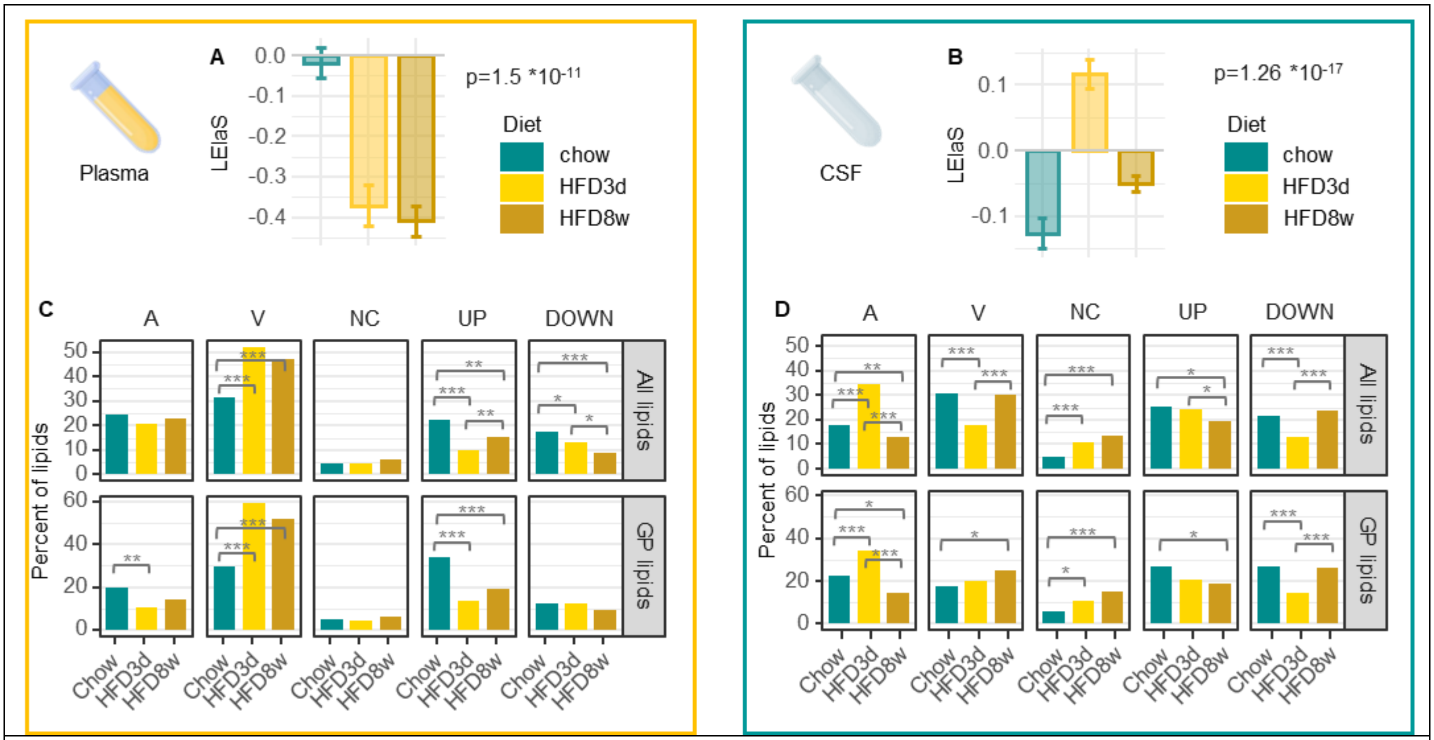


**Supplemental Figure 7**: **Effect of HFD on the elasticity of lipids in Plasma and CSF.**
**A-B** Average signed elasticity scores for all lipids from the brain from mice fed chow, HFD 3d or HFD 8w**,** in plasma (A) and the cerebrospinal fluid (CSF, B).
**C-D** Proportion (in %) of total number of lipid species detected (all, top or glycerophospholipids only, bottom) that belong to each shape group (A, V, NC ("no change"), UP, DOWN), in the plasma (C) and the CSF (D).
Statistics: A-B: Non-adjusted one-way ANOVA p-values comparing diets for each class are displayed on the graphs. C-D: For each tissue, differences in the distribution of lipid-shape categories (A, V, NC, DOWN, UP) across diets were tested using a chi-square test of independence on lipid-species counts (diet × shape). Where indicated, post-hoc pairwise comparisons for each shape between diets were performed using two-proportion tests, with Benjamini–Hochberg correction for multiple testing. p=*0.05' **0.01 ***0.001.

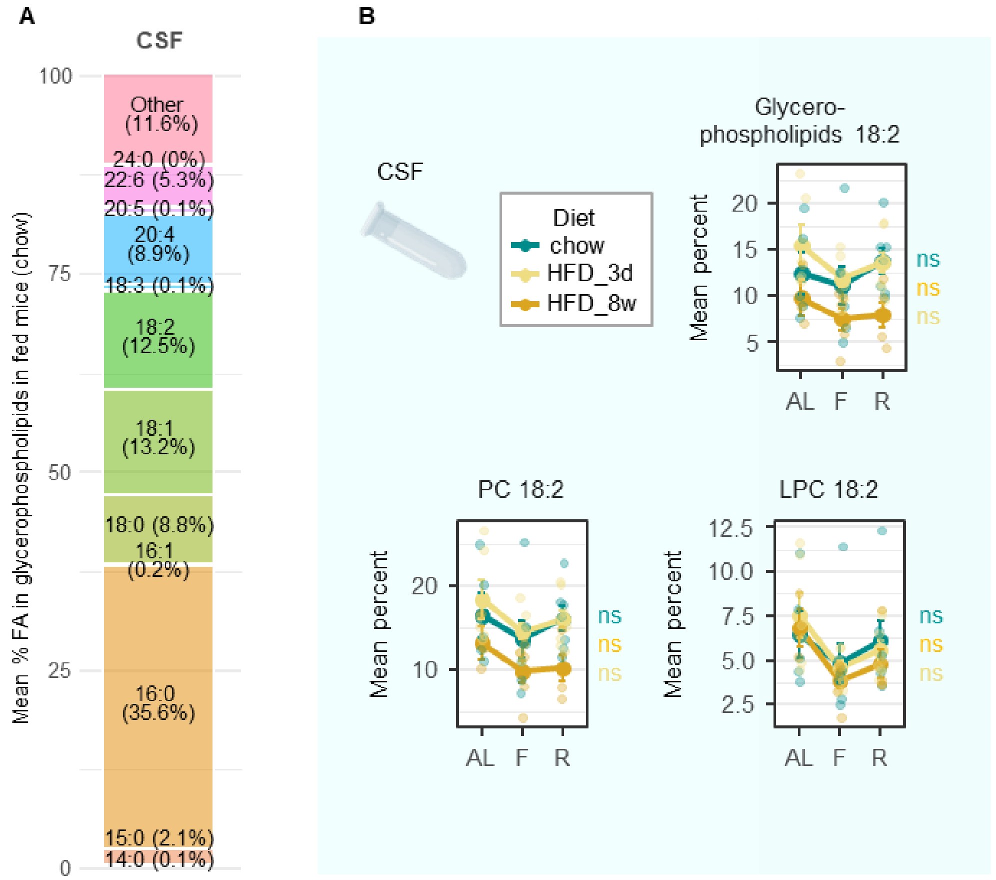


**Supplemental figure 8**: **Phospholipids containing fatty acid 18:2 lose elasticity with high fat diet feeding in cerebrospinal fluid.**
**A** Proportion of glycerophospholipids containing different fatty acids in cerebrospinal fluid (CSF).
**B** Diet-dependent changes in the relative abundance of lipids containing FA 18:2 across feeding states in CSF: glycerophospholipids 18:2 (top right), PC 18:2 (bottom left)**,** LPC 18:2 (bottom right).

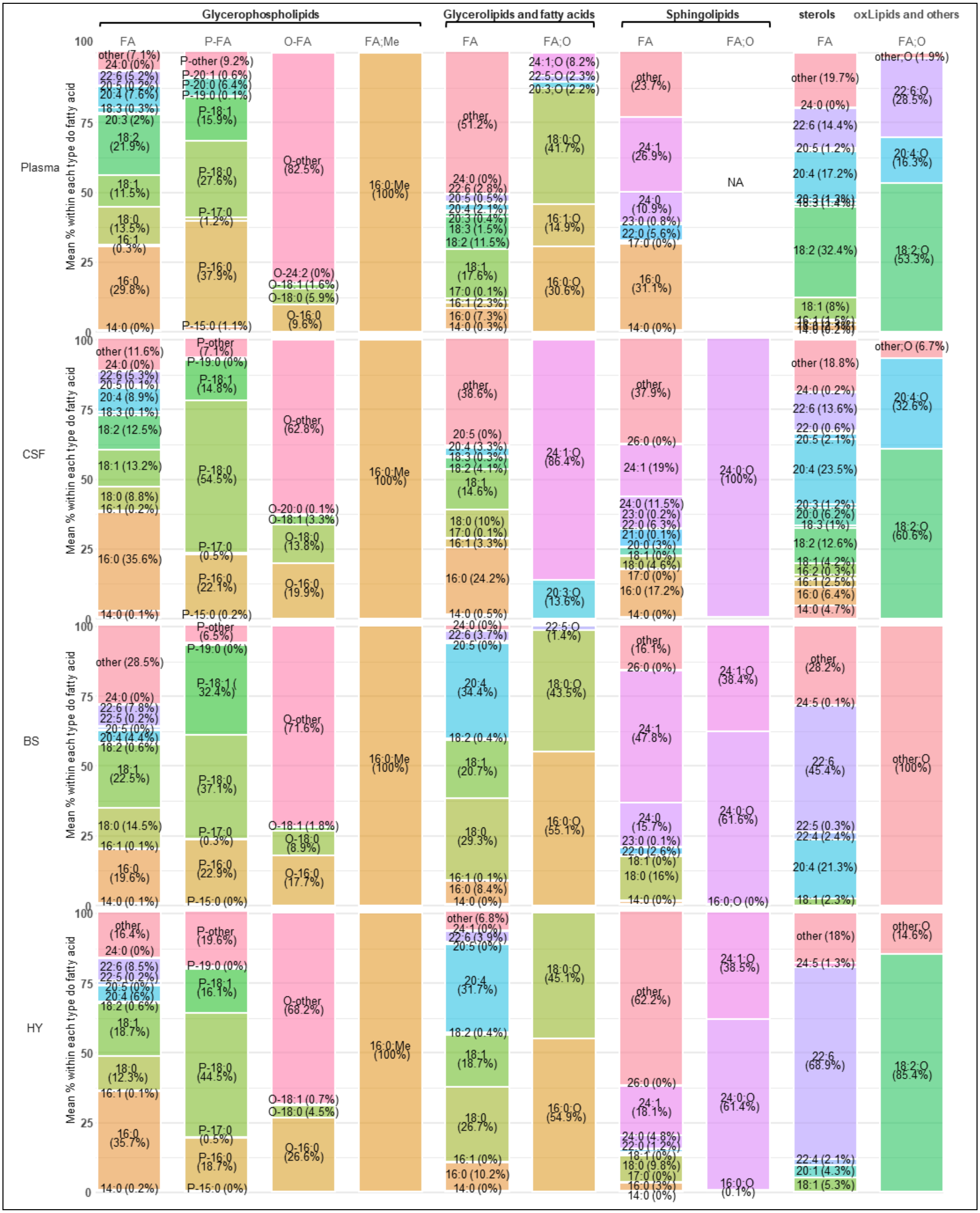
Glycerophospholipids
Glycerolipids and fatty acids
Sphingolipids
sterols
oxLipids and others
FA
P-FA
O-FA
FA;Me
FA
FA;O
FA
FA;O
FA
FA;O
Plasma
CSF
BS
HY
Mean % within each type do fatty acid
100
75
50
25
0
other (7.1%)
24:0 (0%)
22:6 (5.2%)
20:4 (7.6%)
18:3 (0.3%)
20:3 (2%)
18:2 (21.9%)
18:1 (11.5%)
18:0 (13.5%)
16:1 (0.3%)
16:0 (29.8%)
14:0 (0%)
P-other (9.2%)
P-20:1 (0.6%)
P-20:0 (6.4%)
P-19:0 (0.1%)
P-18:1 (15.9%)
P-18:0 (27.6%)
P-17:0 (1.2%)
P-16:0 (37.9%)
P-15:0 (1.1%)
O-other (82.5%)
O-24:2 (0%)
O-18:1 (1.6%)
O-18:0 (5.9%)
O-16:0 (9.6%)
16:0;Me (100%)
other (51.2%)
24:0 (0%)
22:6 (2.8%)
20:5 (0.5%)
20:4 (2.1%)
20:3 (0.4%)
18:3 (1.5%)
18:2 (11.5%)
18:1 (17.6%)
17:0 (0.1%)
16:1 (2.3%)
16:0 (7.3%)
14:0 (0.3%)
24:1;O (8.2%)
22:5;O (2.3%)
20:3;O (2.2%)
18:0;O (41.7%)
16:1;O (14.9%)
16:0;O (30.6%)
other (23.7%)
24:1 (26.9%)
24:0 (10.9%)
23:0 (0.8%)
22:0 (5.6%)
17:0 (0%)
16:0 (31.1%)
14:0 (0%)
NA
other (19.7%)
24:0 (0%)
22:6 (14.4%)
20:5 (1.2%)
20:4 (17.2%)
18:2 (32.4%)
18:1 (8%)
other;O (1.9%)
22:6;O (28.5%)
20:4;O (16.3%)
18:2;O (53.3%)
other (11.6%)
24:0 (0%)
22:6 (5.3%)
20:5 (0.1%)
20:4 (8.9%)
18:3 (0.1%)
18:2 (12.5%)
18:1 (13.2%)
18:0 (8.8%)
16:1 (0.2%)
16:0 (35.6%)
14:0 (0.1%)
P-other (7.1%)
P-19:0 (0%)
P-18:1 (14.8%)
P-18:0 (54.5%)
P-17:0 (0.5%)
P-16:0 (22.1%)
P-15:0 (0.2%)
O-other (62.8%)
O-20:0 (0.1%)
O-18:1 (3.3%)
O-18:0 (13.8%)
O-16:0 (19.9%)
16:0;Me 100%)
other (38.6%)
20:5 (0%)
20:4 (3.3%)
18:3 (0.3%)
18:2 (4.1%)
18:1 (14.6%)
18:0 (10%)
17:0 (0.1%)
16:1 (3.3%)
16:0 (24.2%)
14:0 (0.5%)
24:1;O (86.4%)
20:3;O (13.6%)
other (37.9%)
26:0 (0%)
24:1 (19%)
24:0 (11.5%)
23:0 (0.2%)
22:0 (6.3%)
21:0 (0.1%)
20:0 (3%)
18:1 (0%)
18:0 (4.6%)
17:0 (0%)
16:0 (17.2%)
14:0 (0%)
24:0;O (100%)
other (18.8%)
24:0 (0.2%)
22:6 (13.6%)
22:0 (0.6%)
20:5 (2.1%)
20:4 (23.5%)
20:3 (1.2%)
20:0 (6.2%)
18:3 (1%)
18:2 (12.6%)
18:1 (4.2%)
16:2 (0.3%)
16:1 (2.5%)
16:0 (6.4%)
14:0 (4.7%)
other;O (6.7%)
20:4;O (32.6%)
18:2;O (60.6%)
other (28.5%)
24:0 (0%)
22:6 (7.8%)
22:5 (0.2%)
20:5 (0%)
20:4 (4.4%)
18:2 (0.6%)
18:1 (22.5%)
18:0 (14.5%)
16:1 (0.1%)
16:0 (19.6%)
14:0 (0.1%)
P-other (6.5%)
P-19:0 (0%)
P-18:1 ( 32.4%)
P-18:0 (37.1%)
P-17:0 (0.3%)
P-16:0 (22.9%)
P-15:0 (0%)
O-other (71.6%)
O-18:1 (1.8%)
O-18:0 (8.9%)
O-16:0 (17.7%)
16:0;Me (100%)
24:0 (0%)
22:6 (3.7%)
20:5 (0%)
20:4 (34.4%)
18:2 (0.4%)
18:1 (20.7%)
18:0 (29.3%)
16:1 (0.1%)
16:0 (8.4%)
14:0 (0%)
22:5;O (1.4%)
18:0;O (43.5%)
16:0;O (55.1%)
other (16.1%)
26:0 (0%)
24:1 (47.8%)
24:0 (15.7%)
23:0 (0.1%)
22:0 (2.6%)
18:1 (0%)
18:0 (16%)
14:0 (0%)
24:1;O (38.4%)
24:0;O (61.6%)
16:0;O (0%)
other (28.2%)
24:5 (0.1%)
22:6 (45.4%)
22:5 (0.3%)
22:4 (2.4%)
20:4 (21.3%)
18:1 (2.3%)
other;O (100%)
other (16.4%)
24:0 (0%)
22:6 (8.5%)
22:5 (0.2%)
20:5 (0%)
20:4 (6%)
18:2 (0.6%)
18:1 (18.7%)
18:0 (12.3%)
16:1 (0.1%)
16:0 (35.7%)
14:0 (0.2%)
P-other (19.6%)
P-19:0 (0%)
P-18:1 (16.1%)
P-18:0 (44.5%)
P-17:0 (0.5%)
P-16:0 (18.7%)
P-15:0 (0%)
O-other (68.2%)
O-18:1 (0.7%)
O-18:0 (4.5%)
O-16:0 (26.6%)
16:0;Me (100%)
other (6.8%)
24:1 (0%)
22:6 (3.9%)
20:5 (0%)
20:4 (31.7%)
18:2 (0.4%)
18:1 (18.7%)
18:0 (26.7%)
16:1 (0%)
16:0 (10.2%)
14:0 (0%)
18:0;O (45.1%)
16:0;O (54.9%)
other (62.2%)
26:0 (0%)
24:1 (18.1%)
24:0 (4.8%)
22:0 (1.2%)
18:1 (0%)
18:0 (9.8%)
17:0 (0%)
16:0 (3%)
14:0 (0%)
24:1;O (38.5%)
24:0;O (61.4%)
16:0;O (0.1%)
other (18%)
24:5 (1.3%)
22:6 (68.9%)
22:4 (2.1%)
20:1 (4.3%)
18:1 (5.3%)
other;O (14.6%)
18:2;O (85.4%)

**Supplemental figure 9**: **Fatty acid composition by tissue in mice fed chow for different functional groups and fatty acid type**
Proportion of lipids containing different fatty acyl chains in plasma, cerebrospinal fluid (CSF), brainstem (BS) and hypothalamus (HY), by functional group (Glycerophospholipids, Glycerolipids and fatty acids, sphingolipids, sterols and oxylipids/others), further grouped by fatty acyl modification where applicable (unmodified ("FA"), O-linked (O-FA), P-linked (P-FA), Oxydized (FA;O) and methylated (FA;Me).

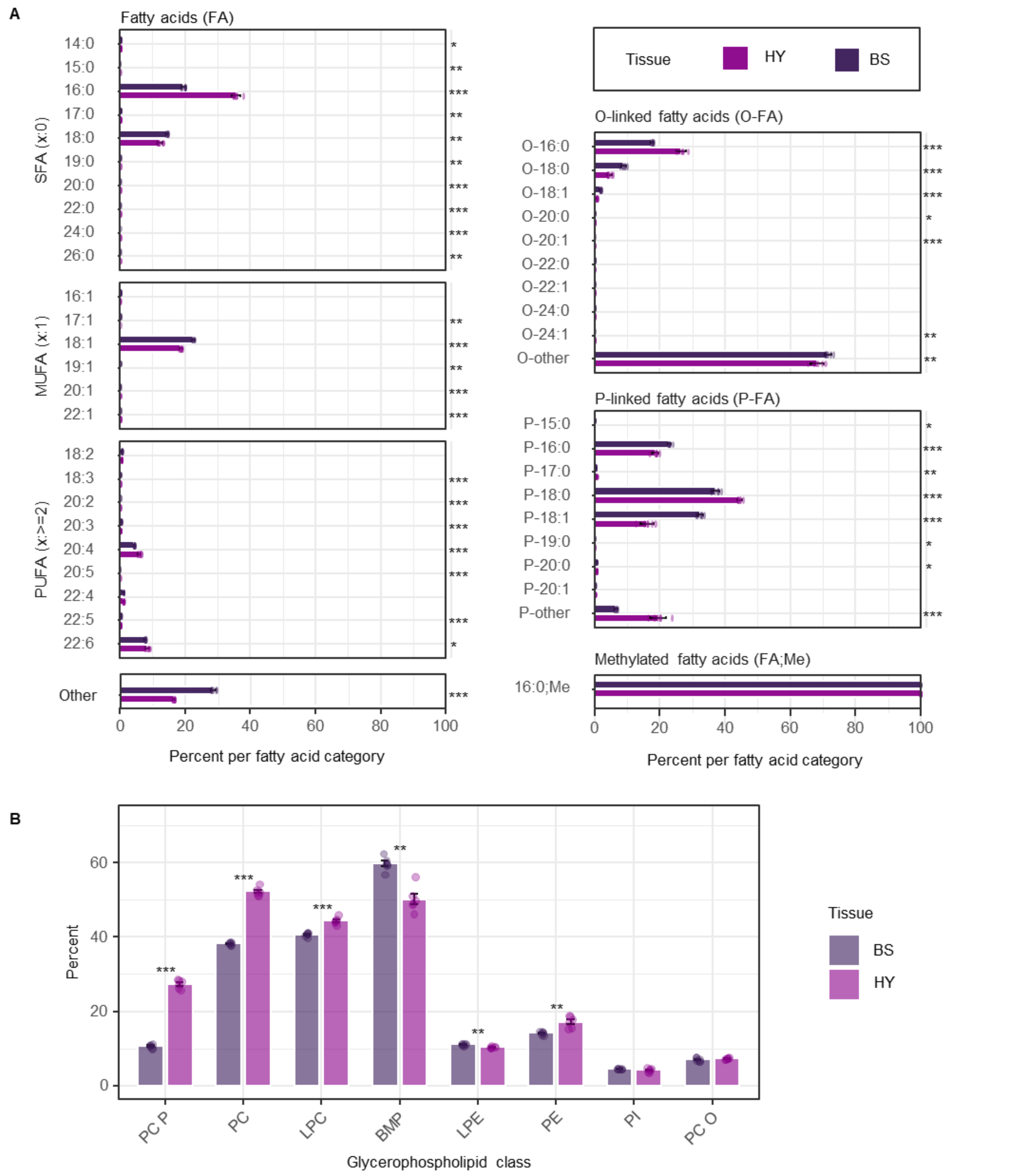


**Supplemental figure 10**: **Comparison of % abundance of lipids containing specific fatty acids in the hypothalamus and brainstem.**

**A** Mean percent of specific fatty acid-containing Glycerophospholipids in the hypothalamus (HY) and brainstem (BS) of mice fed chow ad libitum. SFA: Saturated fatty acids; MUFA: Monounsaturated fatty acids; PUFA: Polyunsaturated fatty acids; other: lipids with 2 or more fatty acids that were not distinguishable in the quantification process. Ether-linked fatty acids: O-FA: O-linked (alkyl); P-FA: P-linked (alkenyl/plasmalogen) fatty acids. FA;Me: methylated fatty acids.

**B** Percent abundance of 16:0 across glycerophospholipid classes in chow fed mice, comparing HY to BS. Each dot represents an individual mouse. Sample size, n = 6 mice/ tissue. Statistical tests: Student t-test +BH correction. *0.05, **0.01, ***0.001

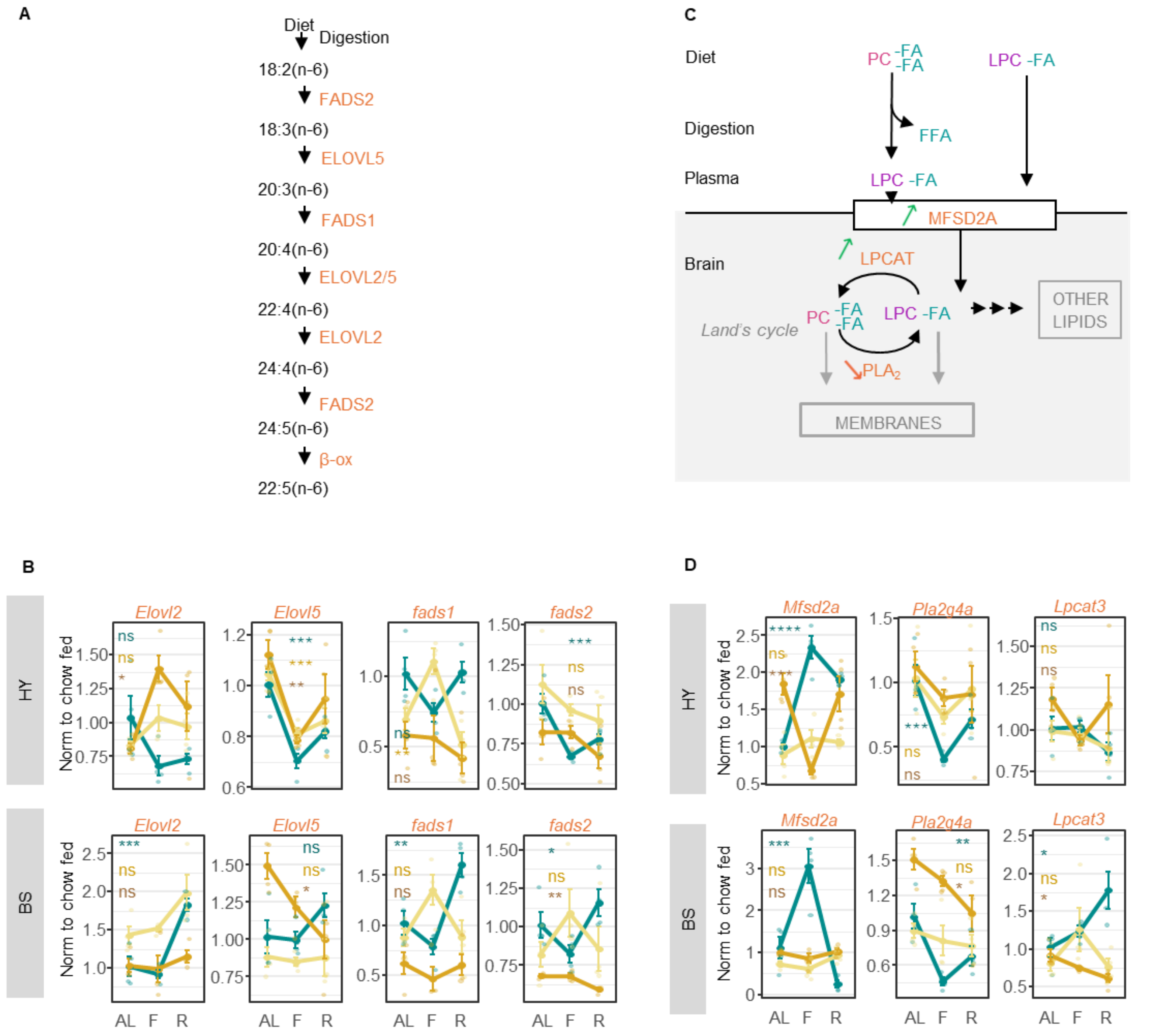


**Supplemental figure 11: Effect of high fat diet on the elasticity of genes involved in 18:2-containing lipids metabolism.**

**A** Schematic of molecular mechanism of enzymes involved in essential fatty acid metabolism diagram adapted from Kawaguchi et al (Comprehensive Natural Products Chemistry, 1999).

**B** Changes in expression of genes involved in the processing of de novo fatty acid synthesis and essential fatty acid metabolism in the hypothalamus (HY, top) and brainstem (BS, bottom) of mice fed different diets under different feeding statuses. Values are normalized to chow fed group.

**C** Schematic of molecular mechanism for of PC import from plasma to the brain. PC and LPC from the diet are digested and absorbed into circulation, however only LPCs can pass the blood brain barrier through the action of the transporter MFSD2A. Once in the brain, LPCs can be transformed back into PCs through the land's cycle or converted to other lipids. Both PCs and LPCs can be incorporated into membranes. Coloured arrows indicate changes in expression with fasting in mice fed chow as shown in H (green= increase, red = decrease). diagram adapted from Semba et al (Advances in Nutrition, 2020)

**D** Changes in expression of key hypothalamic enzymes responsible for LPC transport across the blood brain barrier (Mfsd2a) and enzymes from the land's cycle (Pla2g4a, Lpcat3) metabolism in the hypothalamus (HY, top) and brainstem (BS, bottom) of mice fed different diets under different feeding statuses. Values are normalized to chow fed group.
Feeding status: ad libitum fed (AL), fasted (F), refed (R).
Diets: chow (cyan), HFD 3d (yellow) and HFD 8w (orange).
Data are represented as mean ± SEM; individual mouse values are overlaid. N= 6-7 mice / feeding status/ diet; One way ANOVA comparing status within each diet; p=*0.05' **0.01 ***0.001.
